# Pharmacologic Activation of a Compensatory Integrated Stress Response Kinase Promotes Mitochondrial Remodeling in PERK-deficient Cells

**DOI:** 10.1101/2023.03.11.532186

**Authors:** Valerie Perea, Kelsey R. Baron, Vivian Dolina, Giovanni Aviles, Jessica D. Rosarda, Xiaoyan Guo, Martin Kampmann, R. Luke Wiseman

**Author notes:** Corresponding Author: R. Luke Wiseman, Department of Molecular Medicine The Scripps Research Institute, La Jolla, CA 92037. Authors contributed equally (Valerie Perea, Kelsey R. Baron).

## Abstract

The integrated stress response (ISR) comprises the eIF2α kinases PERK, GCN2, HRI, and PKR, which induce translational and transcriptional signaling in response to diverse insults. Deficiencies in PERK signaling lead to mitochondrial dysfunction and contribute to the pathogenesis of numerous diseases. We define the potential for pharmacologic activation of compensatory eIF2α kinases to rescue ISR signaling and promote mitochondrial adaptation in PERK-deficient cells. We show that the HRI activator BtdCPU and GCN2 activator halofuginone promote ISR signaling and rescue ER stress sensitivity in PERK-deficient cells. However, BtdCPU induces mitochondrial depolarization, leading to mitochondrial fragmentation and activation of the OMA1-DELE1-HRI signaling axis. In contrast, halofuginone promotes mitochondrial elongation and adaptive mitochondrial respiration, mimicking regulation induced by PERK. This shows halofuginone can compensate for deficiencies in PERK signaling and promote adaptive mitochondrial remodeling, highlighting the potential for pharmacologic ISR activation to mitigate mitochondrial dysfunction and motivating the pursuit of highly-selective ISR activators.

## INTRODUCTION

Endoplasmic reticulum (ER) stress and mitochondrial dysfunction are inextricably linked in the onset and pathogenesis of etiologically diverse diseases including cancer, diabetes, and many neurodegenerative disorders. This has led to considerable interest in defining the biological mechanisms responsible for regulating mitochondria during ER stress. The PERK arm of the unfolded protein response (UPR) has emerged as an important stress-responsive signaling pathway for adapting mitochondria in response to pathologic ER insults (Almeida *et al*, 2022; Rainbolt *et al*, 2014). PERK is an ER-localized kinase that is activated in response to ER stress through a mechanism involving autophosphorylation and dimerization (Gardner *et al*, 2013; Hetz *et al*, 2020; Walter & Ron, 2011). Once activated, PERK primarily functions through selective phosphorylation of the α subunit of eukaryotic initiation factor 2 (eIF2α). This leads to both a reduction in new protein synthesis and the selective activation of stress-responsive transcription factors such as ATF4 that regulate expression of genes involved in many adaptive pathways including cellular redox, amino acid biosynthesis, and cellular proteostasis (Han *et al*, 2013; Harding *et al*, 2000; Wek & Cavener, 2007).

PERK localizes to ER-mitochondrial contact sites, positioning this ER stress sensor to coordinate ER and mitochondria function in response to pathologic insults (Munoz *et al*, 2013; Verfaillie *et al*, 2012). Consistent with this, PERK-regulated transcriptional and translational signaling are both implicated in the adaptive remodeling of mitochondrial morphology and function (Almeida *et al*., 2022; Rainbolt *et al*., 2014). PERK signaling promotes both protective mitochondrial elongation and mitochondrial cristae formation in response to ER stress through mechanisms including adaptive remodeling of mitochondrial phospholipids and regulated import of the MICOS subunit MIC19 (Barad *et al*, 2022; Latorre-Muro *et al*, 2021; Lebeau *et al*, 2018; Perea *et al*, 2022). This organellar and ultrastructural remodeling functions to regulate mitochondrial bioenergetics and prevent premature mitochondrial fragmentation during ER stress. Similarly, ATF4-dependent upregulation of SCAF1 downstream of PERK increases assembly of respiratory chain supercomplexes to further adapt mitochondrial bioenergetics during ER stress (Balsa *et al*, 2019). PERK signaling also regulates mitochondrial proteostasis through multiple mechanisms including ATF4-dependent expression of mitochondrial proteostasis factors (e.g., *HSPA9*, *LON*) and reductions in the core TIM23 import subunit TIM17A downstream of translation attenuation (Han *et al*., 2013; Harding *et al*, 2003; Hori *et al*, 2002; Rainbolt *et al*, 2013). Apart from these adaptive functions, PERK signaling regulates mitochondria-derived apoptotic signaling following prolonged, severe ER stress through multiple mechanisms primarily involving sustained upregulation of the transcription factor CHOP (Hetz & Papa, 2018). Thus, PERK serves a central role in dictating mitochondrial adaptation and cellular survival in response to varying levels of ER stress.

Deficiencies in PERK activity induced by genetic, environmental, or aging-related factors are implicated in the onset and pathogenesis of numerous diseases (Almeida *et al*., 2022). Loss-of-function mutations in *EIF2AK3*, the gene that encodes PERK, are causatively associated with Wolcott-Rallison syndrome – a rare autosomal-recessive disorder that involves multi-organ failures including prominent neonatal or early-childhood insulin-dependent diabetes, kidney and liver dysfunction, and cardiac abnormalities (Delepine *et al*, 2000; Julier & Nicolino, 2010; Mann *et al*, 2022). Hypomorphic *EIF2AK3* alleles are also implicated in neurodegenerative diseases including the tauopathy progressive supranuclear palsy (PSP) (Hoglinger *et al*, 2011; Park *et al*, 2022; Yuan *et al*, 2018). Further, deficiencies in PERK signaling have been implicated in the pathogenesis of many diseases including PSP and Huntington’s disease (HD) (Almeida *et al*., 2022; Bruch *et al*, 2017; Ganz *et al*, 2020; Shacham *et al*, 2021). Intriguingly, the pathogenesis of these diseases all involves mitochondrial dysfunction, suggesting that impaired PERK signaling could contribute to disease pathology through deficient regulation of mitochondria.

The above results suggest that pharmacologic enhancement of PERK signaling offers a potential opportunity to mitigate pathologic cellular and mitochondrial dysfunction caused by PERK deficiency. Multiple compounds have been identified to activate PERK signaling including CCT020312 and MK28 (Ganz *et al*., 2020; Grandjean & Wiseman, 2020; Stockwell *et al*, 2012). Both of these compounds have been shown to be beneficial in cellular and mouse models of neurodegenerative diseases including PSP and HD (Bruch *et al*., 2017; Ganz *et al*., 2020). However, the mechanism by which these compounds activate PERK, their selectivity for PERK signaling over other stress-responsive signaling pathways, and the specific dependence of the observed protection on PERK activity remain poorly defined. Further, PERK activating compounds are likely to be limited in their ability to promote protective PERK-dependent signaling in cells expressing inactive or hypomorphic PERK variants. Thus, different strategies are likely required to fully access the therapeutic potential for enhancing adaptive PERK signaling in the context of disease.

An alternative strategy to promote cellular and mitochondria remodeling in PERK-deficient cells is to activate compensatory kinases that induce similar signaling to that regulated by PERK. The integrated stress response (ISR) comprises four stress-activated eIF2α kinases activated in response to diverse pathologic insults (Costa-Mattioli & Walter, 2020; Pakos-Zebrucka *et al*, 2016) (**Fig. 1A**). Apart from PERK, these include GCN2 (activated by amino acid deprivation), HRI (activated by heme deficiency, oxidative stress, and mitochondrial dysfunction), and PKR (activated by double stranded RNA). The activation of these kinases leads to eIF2α phosphorylation and similar translational and transcriptional signaling to that observed upon PERK activation. This suggests that pharmacologic activation of these other ISR kinases could compensate for deficiencies in PERK signaling and rescue pathologic cellular and mitochondrial dysfunction induced by reduced PERK activity. Consistent with this idea, HRI is activated in *Perk*-deficient neurons, but not *Perk*-deficient astrocytes, to promote translational attenuation and ATF4 activation downstream of eIF2α during ER stress (Wolzak *et al*, 2022). While this may reflect mitochondrial stress-dependent activation of HRI signaling induced by ER stress in neurons lacking *Perk* (Fessler *et al*, 2020; Guo *et al*, 2020), these results support the potential for pharmacologically enhancing alternative eIF2α kinases to promote ISR signaling in *Perk*-deficient cells.

**Figure 1.**
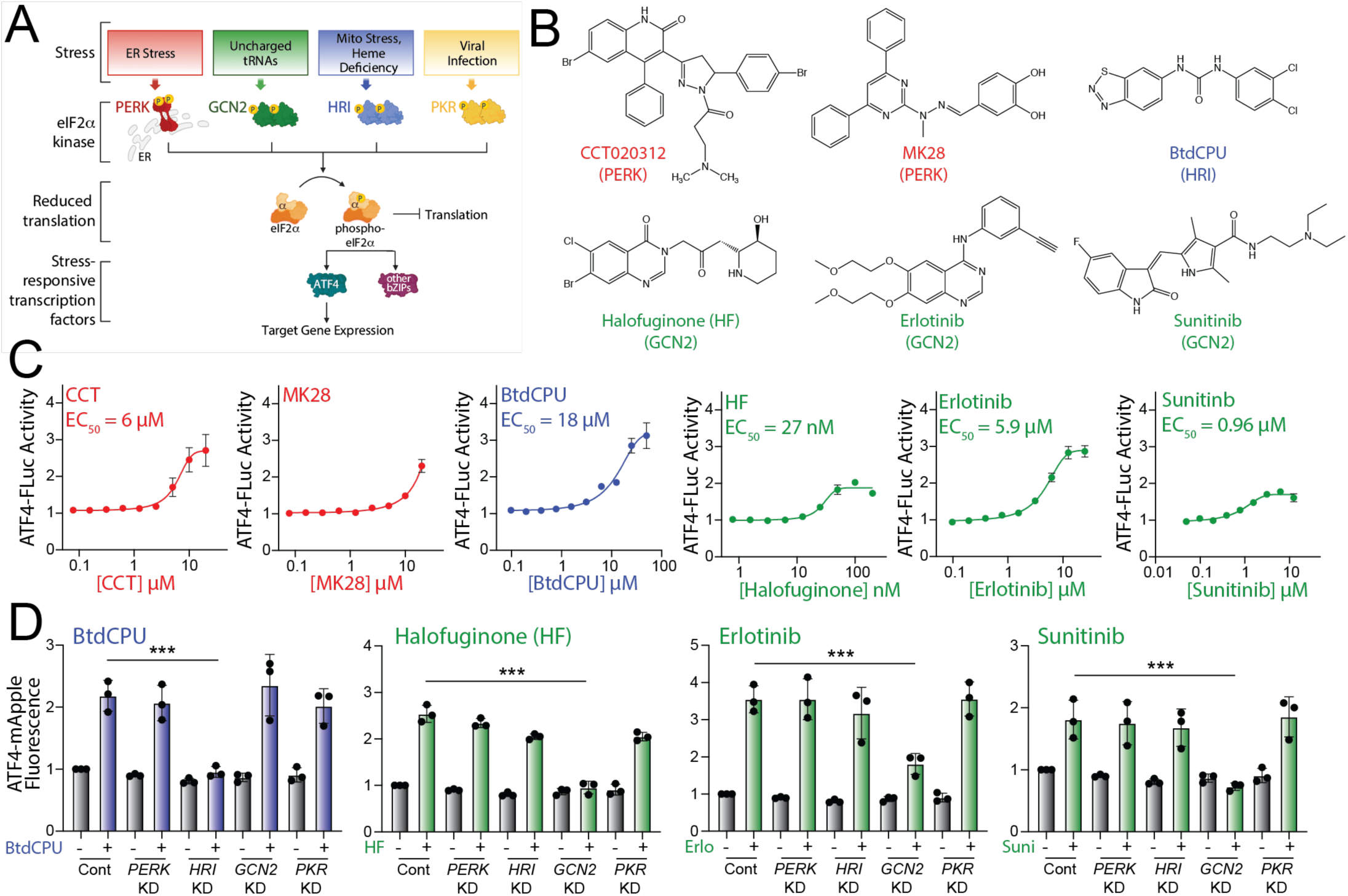
Pharmacologic activation of integrated stress response (ISR) kinases. **A.** Illustration showing the integrated stress response (ISR) comprising the stress-responsive eIF2α kinases PERK, GCN2, HRI, and PKR. Specific cellular insults that activate each individual kinase are shown. **B.** Structures of putative small molecule activators of PERK, HRI, or GCN2 used in this study. **C.** Graphs showing dose-responsive activation of an ATF4-FLuc ISR reporter (**Fig. S1A**) stably expressed in HEK293T cells treated for 8 h with the indicated ISR activator. The EC_50_ of activation is shown. Error bars show SEM for n=12 replicates. **D**. Graphs showing activation of an ATF4-mApple ISR reporter (**Fig. S1A**) stably expressed in HEK293T cells CRISPRi-depleted of the indicated ISR kinase and treated for 8 h with BtdCPU (10 µM), halofuginone (100 nM), Erlotinib (25 µM), or Sunitinib (10 µM). Error bars show SEM for n=3 replicates. ***p<0.005 for one-way ANOVA.

Numerous small molecules have been identified to activate these other ISR kinases. BtdCPU and related N,N’-diarylureas are activators of HRI signaling, although the mechanism by which these compounds activate HRI are poorly defined (Chen *et al*, 2011; Zhang *et al*, 2020). Alternatively, GCN2 is activated by the glutamyl-prolyl tRNA synthetase inhibitor halofuginone through a mechanism involving the accumulation of uncharged proline tRNA (Keller *et al*, 2012). Tyrosine kinase inhibitors such as erlotinib and sunitinib were also shown to activate GCN2 through a mechanism potentially involving direct binding to this ISR kinase (Tang *et al*, 2022). Like PERK activators, these other ISR kinase activators, most notably BtdCPU and halofuginone, have been shown to be protective in cellular and in vivo models of numerous disorders, further highlighting the potential for pharmacologic ISR activation in disease (Chen *et al*., 2011; Ishii *et al*, 2009; Juarez *et al*, 2012; Keller *et al*., 2012; Tian *et al*, 2021). However, the capacity for these compounds to promote adaptive mitochondrial remodeling in wild-type cells or cells deficient in PERK activity is currently unknown.

Here, we define the potential for pharmacologic, stress independent activation of ISR kinases to promote adaptive cellular and mitochondrial remodeling in wild-type and *Perk*-deficient cells. We identify two prioritized ISR activators, BtdCPU and halofuginone, that restores ISR signaling and ER stress sensitivity in *Perk*-deficient cells. However, we find that these compounds differentially impact mitochondria. We show that BtdCPU induces mitochondrial uncoupling, which in turn leads to mitochondrial fragmentation and ISR activation through the OMA1-DELE1-HRI mitochondrial stress signaling axis (Fessler *et al*., 2020; Guo *et al*., 2020). In contrast, halofuginone induces ISR-dependent mitochondrial elongation and adaptive remodeling of mitochondrial respiratory chain activity, mimicking mitochondrial adaptations induced by PERK signaling. These results demonstrate the potential for pharmacologic, stress-independent activation of compensatory ISR kinases to promote adaptive mitochondrial remodeling in wild-type and PERK-deficient cells, motivating the development of highly-selective ISR kinase activating compounds for the treatment of diseases associated with PERK deficiency and/or mitochondrial dysfunction.

## RESULTS

### Activity profiling of ISR activating compounds

The ISR comprises four eIF2α kinases – PERK, HRI, GCN2, and PKR – that are activated in response to diverse stimuli (Costa-Mattioli & Walter, 2020; Pakos-Zebrucka *et al*., 2016) (**Fig. 1A**). We initially probed the activity of previously reported compounds predicted to activate specific ISR kinases. These include the PERK activators CCT020312 and MK28 (Ganz *et al*., 2020; Stockwell *et al*., 2012), the HRI activator BtdCPU (Chen *et al*., 2011), and the GCN2 activators halofuginone, erlotinib, and sunitinib (Keller *et al*., 2012; Tang *et al*., 2022) (**Fig. 1B**). Initially, we tested the ability of these compounds to activate a luciferase-based reporter of ATF4 translation (ATF4-FLuc) as an indicator of ISR activity in HEK293T cells (**Fig. S1A**) (Yang *et al*, 2022). We confirmed that thapsigargin (Tg), a SERCA inhibitor that activates the ER stress responsive PERK ISR kinase, robustly increases ATF4-FLuc activity (**Fig. S1B**). All tested ISR activating compounds activated the ATF4-FLuc reporter with varying potency and efficacy that was consistent with previous reports (**Fig. 1C**) (Chen *et al*., 2011; Ganz *et al*., 2020; Keller *et al*., 2012; Stockwell *et al*., 2012; Tang *et al*., 2022). Next, to determine the specific kinase responsible for compound-dependent ISR activation, we used HEK293T cells expressing an ATF4-mApple translational reporter of ISR activation (**Fig. S1A**) and CRISPRi-depleted of individual ISR kinases (Guo *et al*., 2020). We confirmed the effectiveness of this approach by showing that Tg-dependent ATF4-mApple activation was inhibited in cells deficient in *PERK,* but not cells deficient in other ISR kinases, confirming that ER stress-dependent ATF4 activation is mediated through PERK (**Fig. S1C**). Treatment with CCT020312 or MK28 for 8 h increases ATF4-mApple fluorescence in all cell lines, indicating that, under these conditions, these two compounds activate the ISR reporter through a mechanism not solely dependent on a single kinase (**Fig. S1C**). *HRI*-depletion selectively blocked BtdCPU-dependent ATF4-mApple fluorescence, demonstrating this compound activates the ISR through HRI (**Fig. 1D**). Similarly, *GCN2*-depletion selectively inhibited ATF4-mApple fluorescence induced by halofuginone, erlotinib, and sunitinib, confirming that these compounds activated ISR signaling through GCN2 (**Fig. 1D**). These results show that BtdCPU, halofuginone, erlotinib, and sunitinib activate the ISR through the activity of specific ISR kinases, confirming previously published results (Chen *et al*., 2011; Keller *et al*., 2012; Tang *et al*., 2022).

### Compensatory ISR kinase activation restores ISR signaling in Perk-deficient cells

We next sought to identify compounds that could restore ISR signaling and mitigate pathologic phenotypes linked to deficiencies in PERK activity. Initially, we treated *Perk^+/+^* and *Perk^-/-^* MEFs with ISR activating compounds for 3 h and monitored expression of the ISR target protein ATF4. Tg-dependent increases of ATF4 were blocked in *Perk^-/-^* MEFs, further confirming that ER stress induces ATF4 through a PERK-regulated mechanism (**Fig. 2A,B**). In contrast, BtdCPU, halofuginone, erlotinib, sunitinib, CCT020312, and MK28 increased ATF4 in both *Perk^+/+^* and *Perk^-/-^* MEFs, although *Perk*-deficient cells did show lower levels of ATF4 induction for many compounds, as compared to wild-type cells (**Fig. 2A,B**). Considering other activities of erlotinib (e.g., EGFR inhibition) and sunitinib (e.g., receptor tyrosine kinase inhibition) and the promiscuity for ISR kinase activation observed for CCT020312 and MK28 (**Fig. S1C**), we prioritized BtdCPU and halofuginone for further study. We showed that BtdCPU and halofuginone did not increase ATF4 in knockin MEFs expressing the non-phosphorylatable eIF2α mutant S51A (MEF^A/A^) (Scheuner *et al*, 2001), confirming that the observed increase in ATF4 afforded by these compounds could be attributed to ISR signaling (**Fig. S2A**).

**Figure 2.**
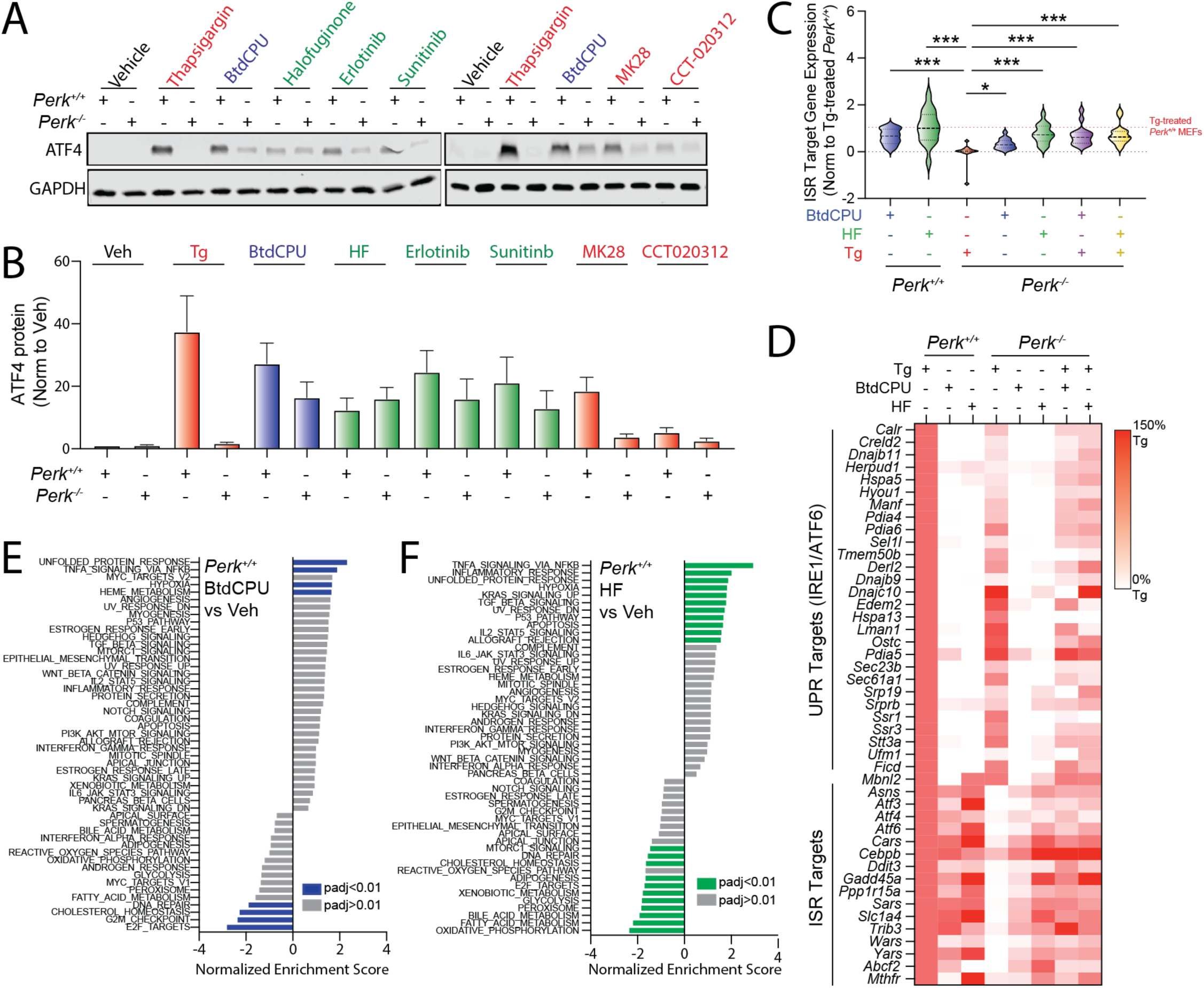
Pharmacologic ISR activators restore ISR signaling in *Perk*-deficient MEFs. **A,B**. Representative immunoblots and quantification of ATF4 in *Perk^+/+^* and *Perk^-/-^*MEFs treated for 3 h with thapsigargin (Tg, 500 nM), BtdCPU (10 µM), halofuginone (100 nM), erlotinib (25 µM), sunitinib (10 µM), MK28 (20 µM), or CCT020312 (10 µM). Error bars show SEM for n=3 replicates. **C.** Expression, measured by RNAseq, of genesets comprised of 16 ISR target genes in *Perk^+/+^* and *Perk^-/-^* MEFs treated for 6 h with thapsigargin (Tg; 500 nM), BtdCPU (10 µM), or halofuginone (HF, 100 nM), as indicated. The expression of individual ISR target genes was normalized to their expression in Tg-treated *Perk^+/+^* MEFs. ISR target genes are described in **Table S2**. *p<0.05, ***p<0.005 for one-way ANOVA relative to Tg-treated *Perk^-/-^* MEFS. **D**. Heat map showing expression, measured by RNAseq, of the UPR and ISR target genes in *Perk^+/+^* and *Perk^-/-^* MEFs treated for 6 h with thapsigargin (Tg; 500 nM), BtdCPU (10 µM), or halofuginone (HF, 100 nM), as indicated. The expression of individual target genes was normalized to their expression in Tg-treated *Perk^+/+^* MEFs. Data are included in **Table S2**. **E,F**. Gene set enrichment analysis (GSEA) for hallmark genesets of RNAseq data from *Perk^+/+^* MEFs treated for 6 h with BtdCPU (10 µM; **E**) or halofuginone (HF, 100 nM; **F**). Full GSEA results are included in **Table S3**.

To further probe the potential for BtdCPU and halofuginone to induce ISR signaling, we monitored gene expression by RNAseq in *Perk^+/+^* and *Perk^-/-^* MEFs treated with these compounds in the presence or absence of Tg (**Table S1**). Initially, we compared the expression of 16 established ISR target genes (Grandjean *et al*, 2019) to that observed in Tg-treated *Perk^+/+^* cells across all conditions (**Fig. 2C,D**, **Table S2**). As expected, *Perk*- deficient cells treated with Tg showed no significant induction of ISR target genes. However, ER stress-responsive genes regulated by the IRE1/XBP1s and ATF6 arms of the UPR are efficiently induced in Tg-treated, *Perk*-deficient cells (**Fig. 2D, Table S2**). Treatment with BtdCPU or halofuginone increased ISR target gene expression in both *Perk^+/+^* and *Perk^-/-^* MEFs in the presence or absence of Tg (**Fig. 2C,D**). We confirmed these results for the ISR target genes *Ddit3/Chop* and *Chac1* by qPCR (**Fig. S2B,C**) These results indicate that BtdCPU and halofuginone both induce ISR signaling in wild-type cells and restore ISR signaling in *Perk*-deficient cells during conditions of ER stress.

Analysis of the transcriptome-wide changes indicate that BtdCPU is a more selective activator of the ISR than halofuginone. Gene set enrichment analysis (GSEA) showed that treatment with BtdCPU (**Table S3**) significantly induced expression of genes related to the unfolded protein response in both *Perk^+/+^* and *Perk^-/-^*MEFs (**Fig. 2E**, **Fig. S2D**). This reflects the increased expression of ISR target genes such as *Atf3, Chac1*, *Asns* and *Atf4* induced by this compound, as markers of other UPR pathways (i.e., IRE1/XBP1s or ATF6) are not significantly induced upon BtdCPU treatment in these cells (**Fig. 2D**). GSEA also identified increased inflammatory and hypoxic signaling in BtdCPU-treated cells (**Fig. 2E**, **Fig. S2D**). However, this appears to be primarily driven by the expression of established ISR target genes such as *Atf3* and *PPP1R15A*, which are included in all these genesets (**Table S3**). By profiling selective transcriptional targets of these and other stress-responsive signaling pathways (Grandjean *et al*., 2019), we found that BtdCPU did not broadly induce activation of other stress-responsive signaling pathways (**Fig. S2E, Table S4**). Thus, our results indicate that BtdCPU is a preferential activator of ISR signaling in these cells.

In contrast, numerous pathways are impacted by halofuginone treatment in *Perk^+/+^* and *Perk^-/-^* MEFs (**Fig. 2F**, **Fig. S2F**). These include the UPR (reflecting ISR activation) and NFκB-mediated inflammatory signaling. Consistent with this, we found that genesets comprised of ISR and NFκB targets are increased by halofuginone treatment in both genotypes (**Fig. S2E, Table S4**). However, halofuginone did not significantly induce expression of genes regulated by other stress-responsive signaling pathways such as the IRE1/XBP1s or ATF6 arms of the UPR, heat shock response (HSR), or oxidative stress response (OSR). Thus, while halofuginone robustly induces stress-responsive signaling through the ISR in both *Perk^+/+^* and *Perk^-/-^* MEFs, our results indicate that this activity is not selective transcriptome-wide.

### BtdCPU and halofuginone reduce ER stress sensitivity of Perk-deficient cells

*Perk-*deficiency leads to increased cellular sensitivity to ER stress (Harding *et al*., 2000). Consistent with this *Perk^-/-^* MEFs show reduced proliferation following Tg treatment, as compared to *Perk^+/+^*MEFs, when measured by crystal violet staining (**Fig. S3A**). Re-overexpression of PERK^WT^ rescues the increased ER stress sensitivity of *Perk^-/-^* MEFs, confirming this effect can be attributed to PERK. Co-treatment with BtdCPU or halofuginone improved proliferation of *Perk*-deficient cells treated with Tg (**Fig. 3A, Fig. S3B**). Other GCN2 activators including erlotinib and sunitinib also showed improved proliferation in Tg-treated *Perk^-/-^* MEFs (**Fig. S3B**). In contast, the putative PERK activator CCT020312 did not increase viability in Tg-treated cells deficient in *Perk*. Further, MK28 showed toxicity in both *Perk^+/+^* and *Perk^-/-^*MEFs, likely reflecting a PERK-independent, off-target activity of this compound.

**Figure 3.**
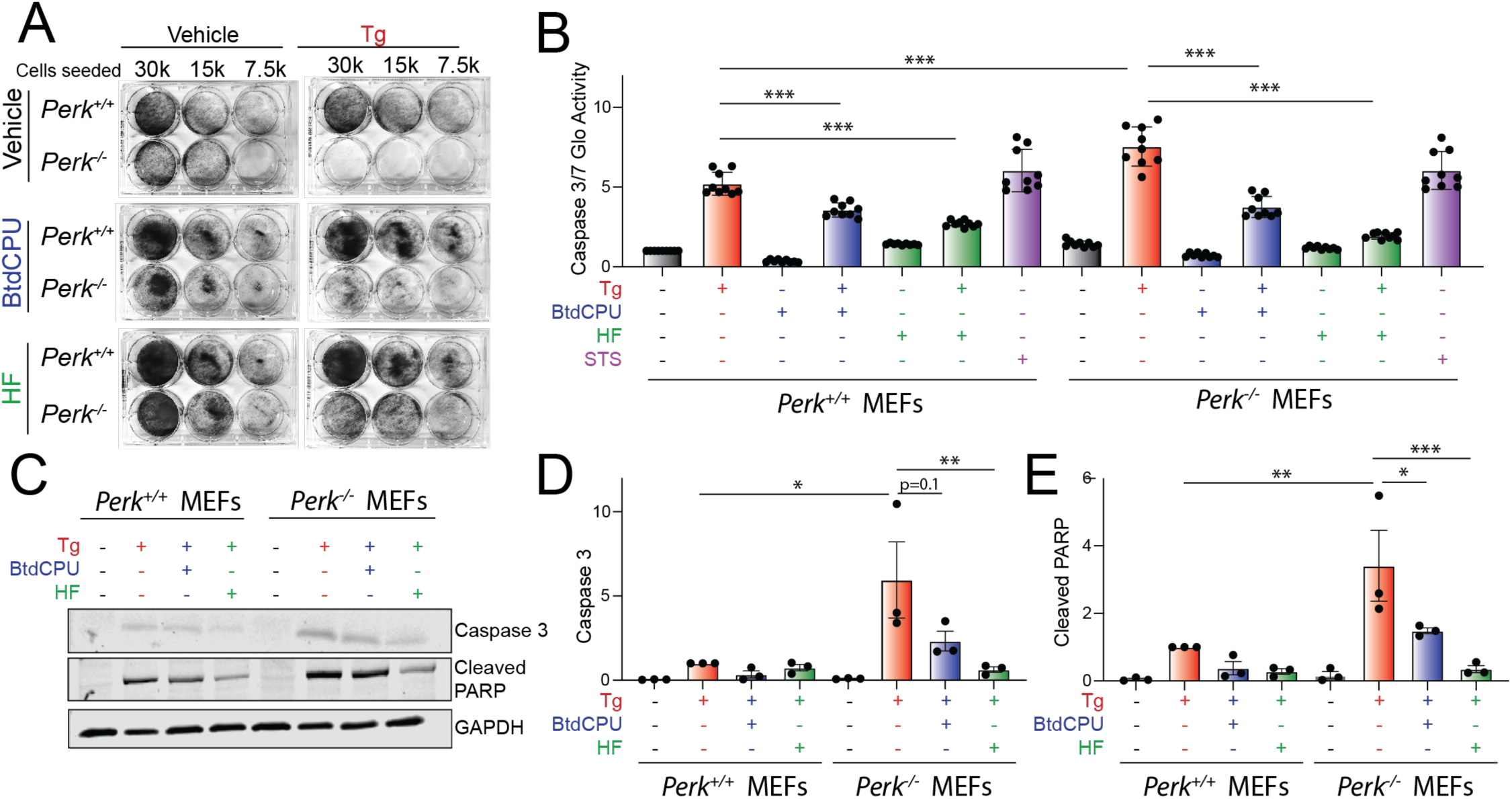
BtdCPU and halofuginone reestablishes ER stress sensitivity in *Perk*-deficient MEFs. **A.** Representative crystal violet staining of *Perk^+/+^* and *Perk^-/-^* MEFs treated for 6 h with thapsigargin (Tg; 500 nM), BtdCPU (10 µM), and/or halofuginone (100 nM) and then replated and allowed to proliferate in 6-well plates. Crystal violet staining was performed 72 h after replating. **B**. Caspase 3/7 activity, measured by Caspase-Glo 3/7, in *Perk^+/+^* and *Perk^-/-^*MEFs treated for 24 h with Tg (500 nM), BtdCPU (10 µM), and/or halofuginone (100 nM). Cells treated with staurosporine (1 µM) (STS) for 24 hr are shown as a control. Error bars show SEM for n=9 replicates. ***p<0.005 for one-way ANOVA. **C-E**. Representative images (**C**) and quantification of active caspase 3 (**D**) or cleaved PARP (**E**) in *Perk^+/+^* and *Perk^-/-^* MEFs treated for 24 h with thapsigargin (Tg; 500 nM), BtdCPU (10 µM), and/or halofuginone (100 nM). Error bars show SEM for n=3. *p<0.05, **p<0.01, ***p<0.005 for one-way ANOVA.

We next monitored the activity of the pro-apoptotic caspases 3 and 7 in *Perk^+/+^* and *Perk^-/-^* MEFs co-treated with Tg and the ISR activators BtdCPU or halofuginone using Caspase-Glo 3/7. Tg-treated *Perk^-/-^* MEFs showed higher caspase 3/7 activity, as compared to Tg-treated *Perk^+/+^* MEFs (**Fig. 3B**). Treatment with BtdCPU or halofuginone, on their own, did not significantly influence caspase 3/7 activity in either *Perk^+/+^* or *Perk^-/-^* MEFs. However, co-treatment with Tg and either BtdCPU or halofuginone reduced Tg-dependent caspase activity in both cell types (**Fig. 3B**). Similar results were observed when caspase activity was monitored by immunoblotting for active, cleaved caspase 3 or cleaved PARP, a caspase substrate (**Fig. 3C-E**) These results indicate that pharmacologic ISR activators such as BtdCPU or halofuginone restores ER stress sensitivity in *Perk*-deficient cells.

### BtdCPU and halofuginone differentially impact mitochondrial morphology

PERK-dependent ISR activation promotes protective mitochondrial elongation in response to ER stress (Lebeau *et al*., 2018; Perea *et al*., 2022). Genetic depletion of *Perk* increases mitochondrial fragmentation and ablates ER stress-dependent increases in elongated mitochondria (Lebeau *et al*., 2018; Perea *et al*., 2022). Thus, we sought to determine whether compensatory ISR kinase activation could rescue mitochondrial elongation in *Perk*- deficient cells. We monitored mitochondrial morphology in *Perk^+/+^* and *Perk^-/-^*MEFs transiently expressing mitochondrial-targeted GFP (^mt^GFP) treated with Tg, BtdCPU, or halofuginone. As reported previously (Lebeau *et al*., 2018; Perea *et al*., 2022), *Perk^-/-^*MEFs showed increased basal mitochondrial fragmentation and were refractory to Tg-induced mitochondrial elongation. (**Fig. 4A,B**). Further, the increase in elongated mitochondria observed in Tg-treated *Perk^+/+^* MEFs was inhibited by co-treatment with ISRIB – an ISR inhibitor that binds to eIF2B and desensitizes cells to eIF2α phosphorylation (Schoof *et al*, 2021; Sidrauski *et al*, 2013; Tsai *et al*, 2018; Zyryanova *et al*, 2018). These results further confirm that the mitochondria elongation observed in Tg-treated cells is attributed to PERK-dependent ISR signaling.

**Figure 4.**
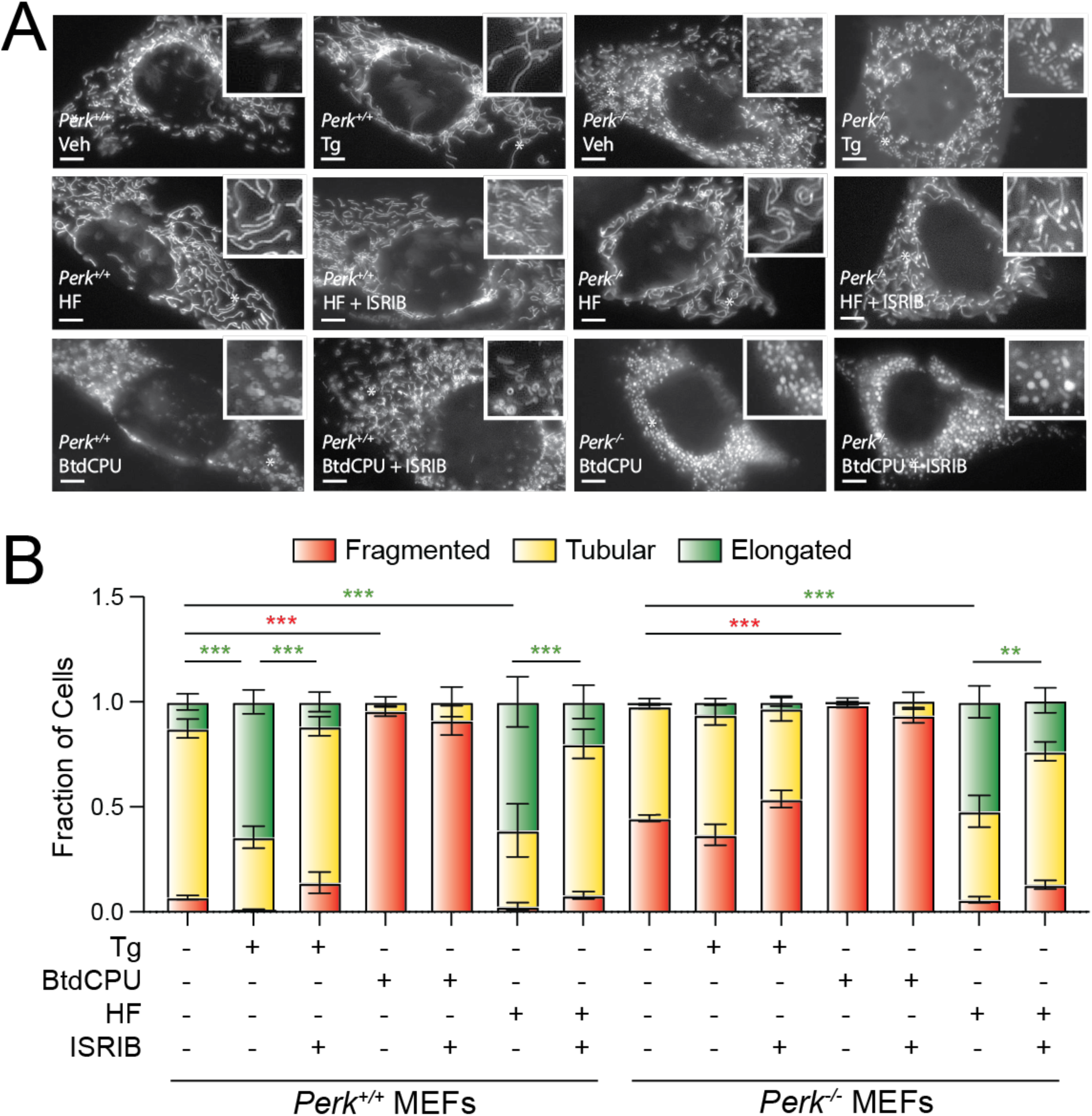
BtdCPU and halofuginone differentially impact mitochondrial morphology. **A,B.**, Representative images and quantification of mitochondrial morphology in *Perk^+/+^* or *Perk^-/-^* MEFs expressing mitochondrial targeted GFP (^mt^GFP) and treated for 3 h with thapsigargin (Tg; 500 nM), BtdCPU (10 µM), halofuginone (HF; 100 nM), and/or ISRIB (200 nM). The inset shows 2-fold magnification of the image centered on the white asterisk. Scale bars, 5 µm. Error bars show SEM for n=3 replicates. *p<0.05, **p<0.01, ***p<0.005 for 2-way ANOVA (red indicates comparison between fragmented mitochondria fractions; green indicates comparisons between elongated mitochondria fractions).

Treatment with BtdCPU did not increase mitochondria elongation (**Fig. 4A,B**). Instead, BtdCPU increased populations of fragmented mitochondria in both *Perk^+/+^* and *Perk^-/-^*MEFs (**Fig. 4A,B**). Similar results were observed in MEF cells stably expressing ^mt^GFP (MEF^mtGFP^; **Fig. S4A**). This increase in fragmentation was not inhibited by co-treatment with ISRIB, indicating that this effect is not attributed to ISR signaling (**Fig. 4A,B**, **Fig. S4A**). In contrast, halofuginone increased mitochondrial elongation in both *Perk^+/+^* and *Perk^-/-^*MEFs to levels similar to that observed in Tg-treated *Perk^+/+^* cells (**Fig. 4A,B**). Halofuginone-dependent increases in elongated mitochondria were also observed in MEF^mtGFP^ cells (**Fig. S4B,C**). Co-treatment with ISRIB inhibited halofuginone-dependent increases in mitochondrial elongation in both genotypes, indicating that the observed elongation results from ISR signaling (**Fig. 4A,B**). Consistent with this, halofuginone treatment demonstrated impaired mitochondrial elongation in MEF^A/A^ cells expressing the non-phosphorylatable S51A eIF2α mutant (Scheuner *et al*., 2001) (**Fig. S4D,E**). These results show that halofuginone and BtdCPU differentially impact mitochondrial morphology. While BtdCPU increases mitochondrial fragmentation, halofuginone promotes adaptive mitochondrial elongation in both *Perk^+/+^*and *Perk^-/-^* MEFs through activation of the ISR, mimicking the mitochondrial elongation induced by PERK activation (Lebeau *et al*., 2018; Perea *et al*., 2022).

### BtdCPU induces mitochondrial depolarization to promote OMA1-DELE1-HRI signaling

Mitochondrial fragmentation is a marker of mitochondrial dysfunction. Thus, the increased fragmentation observed in cells treated with BtdCPU suggests that this compound disrupts mitochondrial activity. To probe this, we monitored mitochondrial respiratory chain activity in *Perk^+/+^* MEFs treated with BtdCPU using Seahorse (**Fig. 5A**). Treatment with BtdCPU did not impact basal respiration, but reduced ATP-linked respiration (**Fig. 5A,B**). This corresponded with an increase of proton leakage, suggesting that BtdCPU impacted mitochondrial membrane potential. Consistent with this, we observed reductions of mitochondrial membrane potential in *Perk^+/+^* MEFs treated with BtdCPU, as measured by TMRE staining (**Fig. 5C**). Similar results were observed in other cell types including HEK293T and SHSY5Y cells (**Fig. S5A,B**). Co-treatment with ISRIB did not influence BtdCPU-dependent mitochondrial depolarization (**Fig. S5B**). This indicates that BtdCPU promotes mitochondrial depolarization independent of ISR activation.

**Figure 5.**
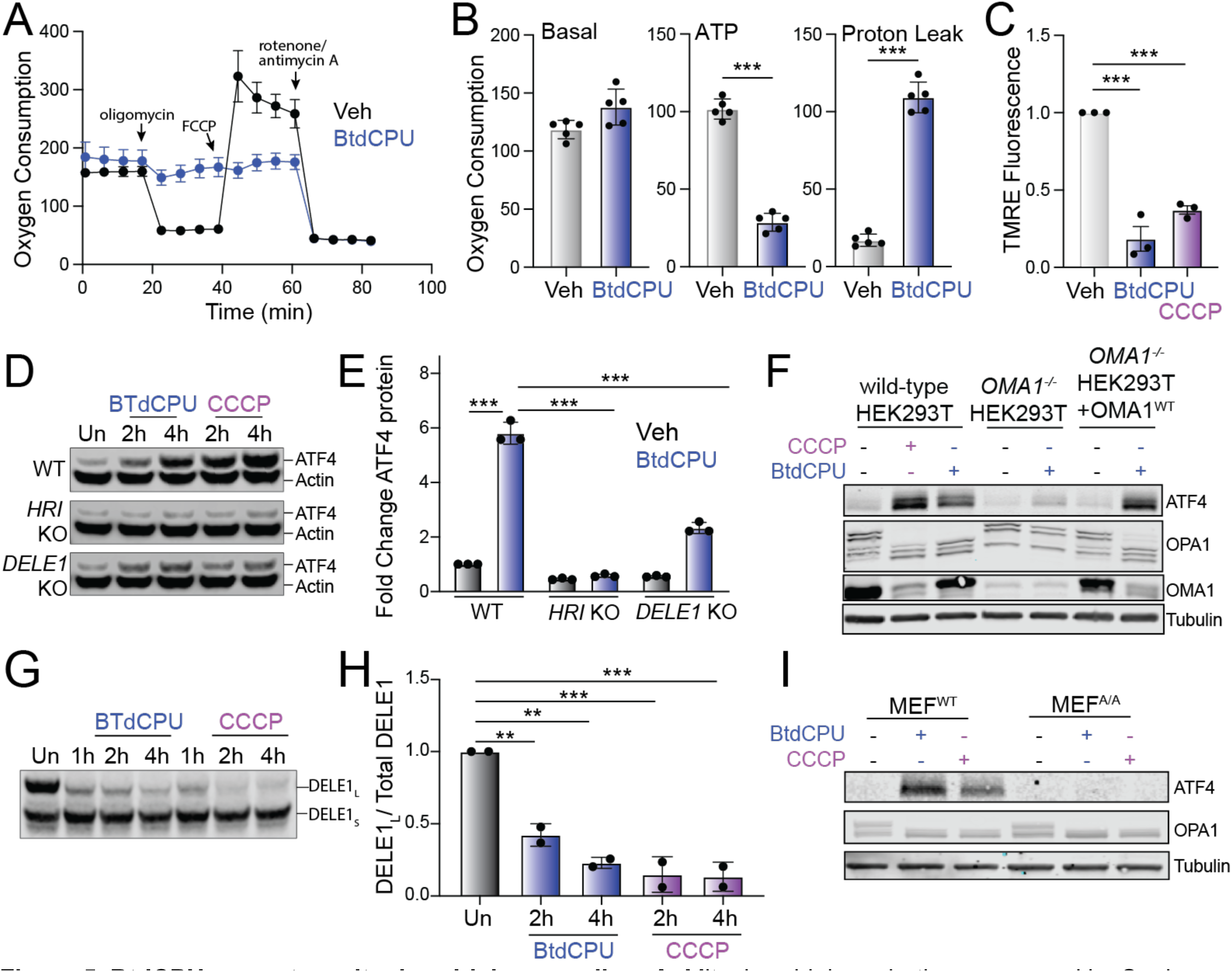
BtdCPU promotes mitochondrial uncoupling. **A.** Mitochondrial respiration, measured by Seahorse mitochondrial stress test, of *Perk^+/+^* MEFs treated for 3 h with vehicle (Veh; black) or BtdCPU (10 µM; blue). Error bars show SEM for n=5 replicates. **B.** Basal respiration, ATP-linked respiration, and proton leak measured from the mitochondrial stress test shown in (**A**). Error bars show SEM for n=5 replicates. *p<0.05, **p<0.01, ***p<0.005 from unpaired t-test. **C**. TMRE fluorescence in *Perk^+/+^*MEFs treated for 3 h with vehicle, BtdCPU (10 µM), or CCCP (25 µM). Error bars show SEM for n=3 replicates. ***p<0.005 from one-way ANOVA. **D,E.** Representative immunoblot (**D**) and quantification (**E**) of ATF4 in wild-type (WT) HEK293T cells or HEK293T cells CRISPR- deleted of *HRI* or *DELE1* treated for the indicated time with BtdCPU (10 µM) or CCCP (5 µM). Actin is showed as a loading control. Error bars show SEM for n=3 replicates. ***p<0.005 for 2-way ANOVA. **F**. Immunoblot of lysates prepared from wild-type HEK293T cells, HEK293T cells CRISPR-deleted of *OMA1*, or HEK293T cells CRISPR-deleted of *OMA1* overexpressing OMA1 treated for 3 h with vehicle, BtdCPU (10 µM), or CCCP (25 µM), as indicated. **G,H**. Representative immunoblot and quantification of long DELE1-mClover (DELE1_L_) cleavage to short DELE1-mClover (DELE1_S_) in HEK293T cells expressing DELE1_L_-mClover treated for the indicated time with BtdCPU (10 µM) or CCCP (5 µM). The cleavage of DELE1_L_ (DELE1_L_ / total DELE1) is shown normalized to untreated cells. Error bars show SEM for n=2 replicates. **p<0.01, ***p<0.005 for one-way ANOVA. **I**. Immunoblot of lysates prepared from wild-type MEFs or MEFs expressing the non-phosphorylatable S51A eIF2α mutant (MEF^A/A^) treated for 3 h with BtdCPU (10 µM) or CCCP (25 µM).

Mitochondrial depolarization can activate ISR signaling through a mechanism involving stress-induced activation of the mitochondrial protease OMA1, cytosolic accumulation of full-length or a C-terminal cleavage products of DELE1, and subsequent DELE1-dependent HRI activation (Fessler *et al*., 2020; Fessler *et al*, 2022; Guo *et al*., 2020). Our results showing that BtdCPU depolarizes mitochondria suggests that this compound could activate ISR signaling through this OMA1-DELE1-HRI signaling axis. Consistent with this, deletion of *HRI* or *DELE1* reduces BtdCPU-dependent ATF4 expression in HEK293T cells, mirroring results observed with the mitochondrial uncoupler CCCP (**Fig. 5D,E**). Similarly, *OMA1-*deletion also inhibited BtdCPU-dependent ATF4 expression (**Fig. 5F**). Re-overexpression of OMA1 restored ATF4 induction in BtdCPU-treated *OMA1-*deficient cells, confirming that this effect can be attributed to OMA1.

OMA1 is a stress-activated protease localized to the inner mitochondrial membrane that is activated in response to mitochondrial stressors such as membrane depolarization (Zhang *et al*, 2014). Once activated, OMA1 induces proteolytic processing of substrates including DELE1 and the inner membrane GTPase OPA1, which induces HRI activation and mitochondrial fission, respectively (Fessler *et al*., 2020; Guo *et al*., 2020; MacVicar & Langer, 2016). We found that BtdCPU increased proteolytic processing of full-length DELE1- mClover expressed in HEK293T cells (**Fig. 5G,H**). Further, OMA1-dependent processing of OPA1 from long to short isoforms was increased in BtdCPU-treated HEK293T cells (**Fig. 5F**). Similar results were observed in MEF^A/A^ cells expressing the non-phosphorylatable S51A eIF2α mutant (Scheuner *et al*., 2001), indicating that this increase of OMA1-dependent OPA1 processing is independent of ISR activity (**Fig. 5I**). We also found that *OMA1*-deletion did not influence BtdCPU-dependent mitochondria depolarization, indicating that OMA1 activation was downstream of mitochondrial uncoupling (**Fig. S5C**). Collectively, these results indicate that BtdCPU-dependent mitochondria depolarization activates OMA1 proteolytic activity to promote mitochondrial fragmentation and ISR activation through a mechanism involving the OMA1-DELE1-HRI signaling pathway, revealing new insights into the mechanism of action for ISR activation afforded by this compound.

### Halofuginone promotes adaptive remodeling of mitochondria respiration in Perk-deficient cells

Mitochondrial elongation is an adaptive mechanism that regulates mitochondrial bioenergetics during conditions of stress (Gomes & Scorrano, 2011; Yao *et al*, 2019). Thus, our results that show PERK signaling impacts mitochondrial morphology both in the absence and presence of acute ER stress suggest an important role for PERK in regulating mitochondrial energy production. To test this, we monitored mitochondrial respiration in *Perk*- deficient MEFs using Seahorse. Cells deficient in *Perk* show increased basal respiration, ATP-linked respiration, spare respiratory capacity, and proton leak, as compared to *Perk^+/+^* MEFs (**Fig 6A,B**). This is in contrast to experiments showing *Perk-*depletion in other cell types including HEK293, myeloid leukemia, and brown adipocyte cells reduce basal respiration (Bassot *et al*, 2023; Grenier *et al*, 2022; Kato *et al*, 2020). However, we demonstrate that re-overexpression of PERK^WT^ in *Perk^-/-^* MEFs restores basal respiratory chain activity to the same levels observed in *Perk^+/+^* MEFs, indicating the observed changes in *Perk*-deficient MEFs can be attributed to the presence of PERK. This likely highlights cell-type specific compensatory regulation of mitochondrial respiration associated with *Perk*-deficiency.

**Figure 6.**
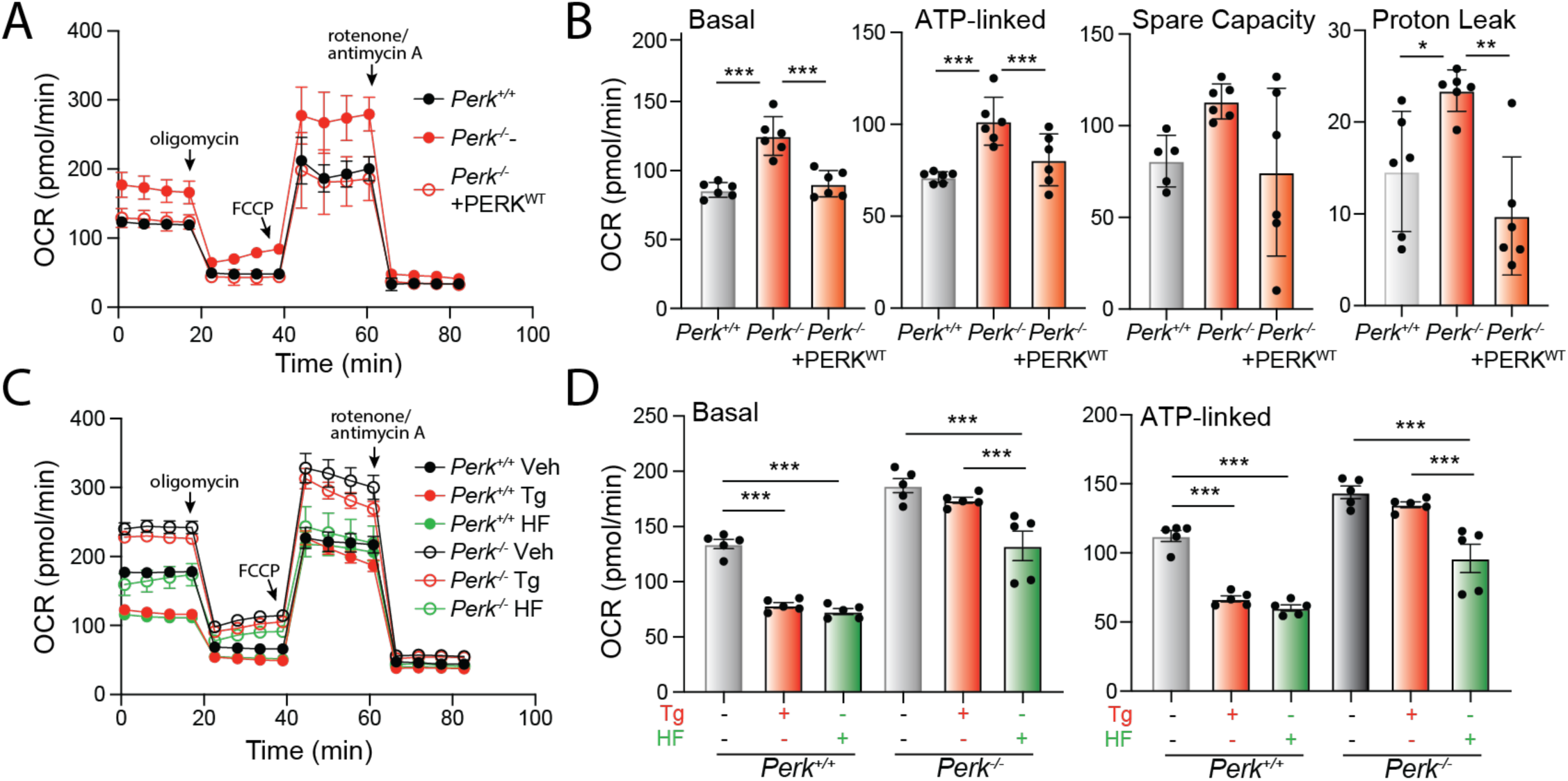
Halofuginone promotes adaptive remodeling of mitochondria respiration in PERK-deficient cells. **A.** Mitochondrial respiration, measured by Seahorse mitochondrial stress test, of *Perk^+/+^*, *Perk^-/-^* MEFs, and *Perk^-/-^* MEFs overexpressing wild-type PERK. Error bars show SEM for n=5 replicates. **B.** Basal respiration, ATP-linked respiration, spare respiratory capacity and proton leak measured from the mitochondrial stress test shown in (**A**). Error bars show SEM for n=5 replicates. *p<0.05, ***p<0.005 for one-way ANOVA **C**. Mitochondrial respiration measured by Seahorse mitochondrial stress test, of *Perk^+/+^*and *Perk^-/-^* MEFs treated for 3 h with thapsigargin (Tg; 500 nM), or halofuginone (HF; 100 nM). Error bars show SEM for n=5 replicates. **D.** Basal respiration and ATP-linked respiration measured from the mitochondrial stress test shown in (**C**). Error bars show SEM for n=5 replicates. ***p<0.005 for one-way ANOVA.

Regardless, we show that *Perk*-deficient MEFs show impaired regulation of respiratory chain activity in response to acute ER stress. In *Perk^+/+^*MEFs, treatment with Tg for 3 h reduces both basal mitochondrial respiration and ATP-linked respiration, while enhancing spare respiratory capacity (**Fig. 6C,D**; **Fig. S6A**). However, these changes were not observed in *Perk*-deficient cells. These results highlight an important role for PERK signaling in regulating mitochondrial respiration in the presence of acute ER stress.

Since halofuginone increases mitochondrial elongation in both *Perk^+/+^* and *Perk^-/-^* MEFs (**Fig. 4A,B**) and mitochondrial elongation is linked to adaptive regulation of mitochondrial respiration (Gomes & Scorrano, 2011; Yao *et al*., 2019), we anticipated that treatment with this compound should promote similar changes to mitochondrial respiration to those observed in wild-type cells treated with PERK-activating ER stressors (e.g., Tg). Consistent with this, treatment of *Perk^+/+^* MEFs with halofuginone induced identical changes of mitochondrial respiration to those observed in Tg-treated cells (**Fig. 6C**). This includes reductions in basal respiration and ATP-linked respiration, and increases in spare respiratory capacity (**Fig. 6D**, **Fig. S6A,B**). Halofuginone also reduced basal respiration and ATP-linked respiration in *Perk^-/-^* MEFs, restoring these parameters to levels similar to those observed in wild-type cells (**Fig. 6C,D**). However, halofuginone did not alter spare respiratory capacity or proton leakage in *Perk*-deficient cells (**Fig. S6A,B**), likely reflecting a chronic consequence of deficient PERK activity on mitochondrial function that cannot be rescued by this short treatment. These results indicate that halofuginone can induce adaptive remodeling of mitochondrial respiration in both wild-type and *Perk*-deficient cells, mimicking changes observed upon PERK activation during ER stress.

## DISCUSSION

Pharmacologic ISR activation has emerged as a promising strategy to mitigate pathologies implicated in etiologically-diverse diseases including many types of cancers, ischemic diseases, and neurodegenerative disorders such as PSP (Almeida *et al*., 2022; Chen *et al*., 2011; Ganz *et al*., 2020; Hughes & Mallucci, 2019; Rosarda *et al*, 2021; Zhang *et al*, 2022). Here, we sought to further define the potential for pharmacologic ISR activation to promote adaptive mitochondrial remodeling and prevent pathologic mitochondrial dysfunction induced under conditions such as *Perk*-deficiency (Almeida *et al*., 2022; Hoglinger *et al*., 2011; Stutzbach *et al*, 2013). By probing the activity of established ISR activating compounds, we demonstrate that treatment with halofuginone restores ER stress sensitivity and promotes adaptive mitochondrial remodeling in both wild-type and *Perk*-deficient cells. These results underscore the potential for pharmacologic ISR kinase activation to mitigate mitochondrial dysfunction associated with diverse disorders.

Numerous compounds have been reported to selectively activate ISR kinases independent of cellular stress; however, each of these compounds has liabilities that limit their ability to probe the biological and therapeutic potential for ISR signaling in cellular and in vivo models of disease. The PERK activator CCT020312 has been used to activate PERK signaling in multiple models, including mouse models of PSP where CCT020312 was shown to promote clearance of tau aggregates (Bruch *et al*., 2017). However, recent results suggest this increased tau clearance reflects an off-target activity of this compound that increases autophagy independent of the ISR, questioning the dependence of this effect on PERK activation (Yoon *et al*, 2022). Similarly, the PERK activator MK28 reduces toxic huntingtin aggregation in mouse models of HD (Ganz *et al*., 2020). However, we found this compound to be cytotoxic in *Perk^+/+^* and *Perk^-/-^* MEFs (**Fig. S3B**), likely reflecting a PERK-independent, off-target activity. Further, the mechanism of both CCT020312- and MK28-dependent PERK activation is currently poorly understood. While elucidation of the mechanism of PERK activation for these compounds could enable the establishment of next generation compounds with improved selectivity and potency, the use of CCT020312 and MK28 as PERK activators should be approached with care and include use of both pharmacologic or genetic controls to confirm that specific phenotypes can be attributed to PERK activation, as opposed to off-target activities.

BtdCPU and related N,N’ diarylureas have largely been developed to increase apoptotic signaling downstream of HRI for the treatment of hematologic cancers including multiple myeloma and acute leukemia (Burwick *et al*, 2017; Smith *et al*, 2021). However, the mechanism of BtdCPU-dependent HRI activation was previously undefined. We demonstrate that BtdCPU disrupts the mitochondrial membrane potential to activate the stress-activated mitochondrial protease OMA1. This leads to ISR activation through the OMA1-DELE1-HRI mitochondrial stress-signaling axis and mitochondrial fragmentation through OMA1-dependent proteolytic processing of OPA1. Since mitochondrial uncoupling is associated many pathologic conditions, this mechanism of action limits the application of BtdCPU and related compounds to promote protective ISR signaling and mitochondrial adaptation in context of other diseases.

Unlike other ISR activator compounds discussed above, the mechanism of halofuginone-dependent ISR activation is known to be attributed to its inhibition of the glutamyl-prolyl tRNA synthetase (Keller *et al*., 2012). This leads to accumulation of uncharged proline tRNAs and subsequent GCN2 activation. Halofuginone has been shown to be protective in models of etiologically-diverse diseases including ischemic disorders, cardiovascular disease, and many cancers (Ishii *et al*., 2009; Juarez *et al*., 2012; Pines & Spector, 2015). However, apart from ISR activation, halofuginone can also promote protection through other mechanisms including inhibition of TGFβ signaling (Pines & Spector, 2015). In addition, our RNAseq transcriptional profiling indicates that halofuginone does not show transcriptome-wide selectivity for the ISR, as it also induces expression of genes regulated by other pathways including NFκB-mediated inflammatory signaling. Further, higher doses of halofuginone can inhibit global translation independent of the ISR, owing to its glutamyl-prolyl tRNA synthetase inhibition (Keller *et al*., 2012; Pitera *et al*, 2022). These effects may limit the translational potential for halofuginone-dependent ISR activation as a strategy to mitigate mitochondrial dysfunction in disease.

Regardless of these limitations, we show that halofuginone-dependent ISR activation both promotes ER stress sensitivity and adaptive mitochondrial remodeling in wild-type and *Perk*-deficient cells. This demonstrates the potential for pharmacologic activation of compensatory ISR kinases to promote adaptive mitochondria remodeling, even in cells deficient in PERK. However, our results also demonstrate the need for continued development of highly selective ISR kinase activators that can be used to further probe the potential for this approach to mitigate cellular and mitochondrial dysfunction in disease. While pharmacologic ISR activation offers unique opportunities to promote protective signaling through this pathway, a challenge for the translational development of pharmacologic ISR kinase activators is the pro-apoptotic signaling that could result from chronic ISR activation (Hetz & Papa, 2018). Previous results have shown that optimization of compound PK/PD can allow transient, *in vivo* activation of similar stress-responsive signaling pathways such as the IRE1/XBP1s signaling arm of the UPR to induce protective, adaptive signaling without the pathologic consequences of chronic pathway activation (Madhavan *et al*, 2022). Similar strategies could be applied for ISR kinase activating compounds to allow protective, adaptive ISR signaling without inducing pro-apoptotic signaling associated with chronic ISR activity. As we, and others, continue identifying ISR kinase activating compounds, the therapeutic potential and optimized activity of this class of compounds will continue to be defined, revealing new insights into the translational potential of pharmacologic ISR kinase activation to mitigate mitochondrial dysfunction for diverse disorders.

## MATERIALS AND METHODS

### Cell Culture, Transfections, Lentiviral Transduction

*Perk^+/+^* and *Perk^-/-^* MEFs (kind gifts from David Ron, Cambridge)(Harding *et al*., 2000) and MEF^A/A^ cells (kind gifts from Randal Kaufman; Sanford-Burnham-Prebys)(Scheuner *et al*., 2001) were cultured as previously described at 37°C and 5% CO_2_ in DMEM (Corning-Cellgro) supplemented with 10% fetal bovine serum (FBS; Omega Scientific), 2 mM L-glutamine (GIBCO), 100 Units/mL penicillin, 100 mg/mL streptomycin (GIBCO), non-essential amino acids (GIBCO), and 2-mercaptoethanol (ThermoFisher). HEK293T cells (purchased from ATCC) and SHSY5Y cells (purchased from ATCC) were cultured at 37°C and 5% CO_2_ in DMEM (Corning-Cellgro) supplemented with 10% fetal bovine serum (FBS; Omega Scientific), 2 mM L-glutamine (GIBCO), 100 Units/mL penicillin, and 100 mg/mL streptomycin (GIBCO). *OMA1*-deficient HEK293T cells, *DELE1*-deficient HEK293T cells, and *HRI*-deficient HEK293T cells were described previously (Guo *et al*., 2020) and cultured at 37°C and 5% CO_2_ in DMEM (Corning-Cellgro) supplemented with 10% fetal bovine serum (FBS; Omega Scientific), 2 mM L-glutamine (GIBCO), 100 Units/mL penicillin, and 100 mg/mL streptomycin (GIBCO). HEK293T cells stably expressing ATF4-FLuc or ATF4-mAPPLE CRISPRi-depleted of *HRI*, *GCN2*, *PERK*, or *PKR* were described previously (Guo *et al*., 2020; Yang *et al*., 2022) and cultured at 37°C and 5% CO_2_ in DMEM (Corning-Cellgro) supplemented with 10% fetal bovine serum (FBS; Omega Scientific), 2 mM L-glutamine (GIBCO), 100 Units/mL penicillin, and 100 mg/mL streptomycin (GIBCO). MEF^mtGFP^ cells (kind gift from Peter Schultz, TSRI)(Wang *et al*, 2012) were cultured at 37°C and 5% CO_2_ in DMEM (Corning-Cellgro) supplemented with 10% fetal bovine serum (FBS; Omega Scientific), 2 mM L-glutamine (GIBCO), 100 Units/mL penicillin, and 100 mg/mL streptomycin (GIBCO). MEF cells were transfected with MEF Avalanche Transfection Reagent (EZ Biosystems) according to the manufacturer’s protocol.

### Plasmids, compounds, and reagents

All compounds used in this study were purchased: thapsigargin (Tg; 50-464-295, Fisher Scientific), ISRIB (SML0843, Sigma), CCCP (C2759, Sigma), BtdCPU (32-489-210MG, Fisher), halofuginone (50-576-30001, Sigma), erlotinib (S7786, Selleckchem), sunitinib (SU11248, Selleckchem), MK-28 (HY-137207, MedChemExpress), CCT020312 (HY-119240, Fisher), staurosporine (S1421, Selleckchem). *Perk^WT^* overexpression plasmid was a kind gift from Jonathan Lin (Stanford) (Yuan *et al*., 2018). The mitochondrial-targeted GFP plasmid (Lebeau *et al*., 2018) and the DELE1_L_-mClover plasmid (Guo *et al*., 2020) were described previously.

### Measurements of ISR activation in ATF4-reporter cell lines

ATF4-FLuc reporter cells were seeded at a density of 15,000 cells per well in Greiner Bio-One CELLSTAR flat 384-well white plates with clear bottoms. The following day, cells were treated with the indicated compound in triplicate 10-point dose response format for eight hours. After treatment, an equal volume of Promega Bright-Glo substrate was added to the wells and allowed to incubate at room temperature for 10 minutes. Luminescence was then measured in an Infinite F200 PRO plate reader (Tecan) with an integration time of 1000 ms.

ATF4-mApple reporter cells were seeded at a density of 300,000 cells per well in 6-well TC-treated flat bottom plates (Genesee Scientific). The following day, cells were treated for eight hours with the indicated concentrations. Following treatment, cells were washed twice with phosphate-buffered saline (PBS) and dissociated using TrypLE Express (Thermo Fisher). The enzymatic reaction was neutralized through addition of flow buffer containing PBS and five percent fetal bovine serum (FBS). Flow cytometry was performed on a Bio-Rad ZE5 Cell Analyzer. mApple (568/592 nm) was measured using the 561 nm green-yellow laser in combination with the 577/15 filter. Analysis was performed using FlowJo^TM^ Software (BD Biosciences).

### Fluorescence Microscopy

MEF or HeLa cells transiently transfected with ^mt^GFP or MEF^mtGFP^ cells were seeded at a density of 100,000 cells/well on glass-bottom dishes (MatTek) coated with poly-D-lysine (Sigma) or rat tail collagen 1 (GIBCO). Cells were then treated as indicated and images were recorded with an Olympus IX71 microscope with 60x oil objective (Olympus), a Hamamatsu C8484 camera (Hamamatsu Photonics), and HCI image software (Hamamatsu Photonics). Quantification was performed by blinding the images and then scoring cells based on the presence of primarily fragmented, tubular, or elongated mitochondria, as before (Lebeau *et al*., 2018). At least three different researchers scored each set of images and these scores were averaged for each individual experiment and all quantifications shown were performed for at least 3 independent experiments quantifying a total of >60 cells/condition across all experiments. The data were then analyzed in PRISM (GraphPad, San Diego, CA) and plotted on a stacked bar plot to show the average morphology and standard error of the mean across all experiments. Statistical comparisons were performed using a 2-way ANOVA in PRISM, comparing the relative amounts of fragmented, tubular, or elongated mitochondria across different conditions.

### Immunoblotting and Antibodies

Whole cells were lysed on ice in HEPES lysis buffer (20 mM Hepes pH 7.4, 100 mM NaCal, 1 mM EDTA, 1% Triton X100) supplemented with 1x Pierce protease inhibitor (ThermoFisher). Total protein concentrations of lysates were then normalized using the Bio-Rad protein assay and lystaes were combined with 1x Laemmli buffer supplemented with 100 mM DTT and boiled for 5 min. Samples (100 µg) were then separated by SDS-PAGE and transferred to nitrocellulose membranes (Bio-Rad). Membranes were blocked with 5% milk in tris-buffered saline (TBS) and then incubated overnight at 4°C with the indicated primary antibody. The next day, membranes were washed in TBS supplemented with Tween, incubated with the species appropriate secondary antibody conjugated to IR-Dye (LICOR Biosciences), and then imaged using an Odyssey Infrared Imaging System (LICOR Biosciences). Quantification was then carried out using the LICOR Imaging Studio software. Primary antibodies were acquired from commercial sources and used in the indicated dilutions in antibody buffer (50mM Tris [pH 7.5], 150mM NaCl supplemented with 5% BSA (w/v) and 0.1% NaN3 (w/v)): ATF4 (Cell Signaling, 1:500), Tubulin [B-5-1-2] (Sigma, 1:5000), HSP60 [LK1] (Thermo Scientific, 1:1000), PERK (C33E10) (Cell Signaling, 1:1000), HA [Clone: 16B12] (Biolegend, 1: 1000), GFP (B2) (Santa Cruz, 1:1000), OPA1 (BD Transduction Labs, 1:2000), OMA1 (Cell Signaling, 1:1000), beta-actin (Cell Signaling 1:5,000), cleaved Caspase 3 (Cell Signaling, 1:1000), PARP (Cell Signaling, 1:1000), GAPDH (Fisher, 1:1000).

### RNA sequencing and analysis

*Perk^+/+^* and *Perk^-/-^* MEFs cells were treated for 6 h with respective compounds as noted. Cells were rinsed with DPBS, lysed, and total RNA was collected using the QuickRNA mini kit (Zymo) according to the manufacturer′s instructions. Transcriptional profiling using whole transcriptome RNA sequencing was conducted via BGI Americas on the DNBseq platform with three biological replicates for each condition. All samples were sequenced to a minimum depth of 20 M PE 150 bp stranded reads. Alignment of reads was performed using DNAstar Lasergene SeqManPro to the mouse genome GRCm39 assembly. Aligned reads were imported into ArrayStar 12.2 with Qseq (DNAStar Inc.) to quantify the gene expression levels. Differential expression analysis and statistical significance comparisons were assessed using DESeq 2 v. 1.34 in R compared to vehicle-treated *Perk^+/+^* cells. Mouse annotations were converted to human orthologs using BiomaRt v. 2.50.3 prior to functional gene set enrichment with the R package fast gene set enrichment (fgsea) v. 1.20.0 using the Hallmark Pathways Geneset v7.5.1 from MsigDB. Code can be provided upon request. The complete RNA-seq data is deposited in gene expression omnibus (GEO) as GSE227134.

### Quantitative Polymerase Chain Reaction (qPCR)

The relative mRNA expression of target genes was measured using quantitative RT-PCR. Cells were treated as indicated and then washed with phosphate buffered saline (PBD; Gibco). RNA was extracted using Quick-RNA MiniPrepKit (Zymo Research) according to the manufacturers protocol. RNA (500 ng) was then converted to cDNA using the QuantiTect Reverse Transcription Kit (Qiagen). qPCR reactions were prepared using Power SYBR Green PCR Master Mix (Applied Biosystems), and primers (below) were obtained from Integrated DNA Technologies. Amplification reactions were run in an ABI 7900HT Fast Real Time PCR machine with an initial melting period of 95 °C for 5 min and then 45 cycles of 10 s at 95 °C, 30 s at 60 °C.

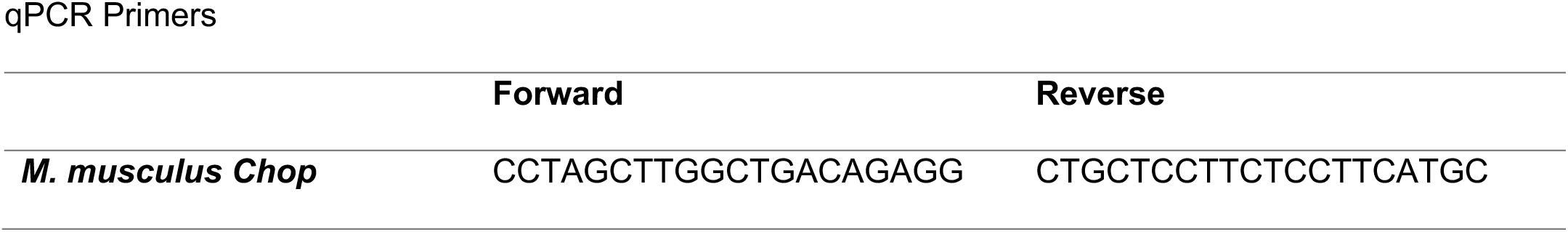
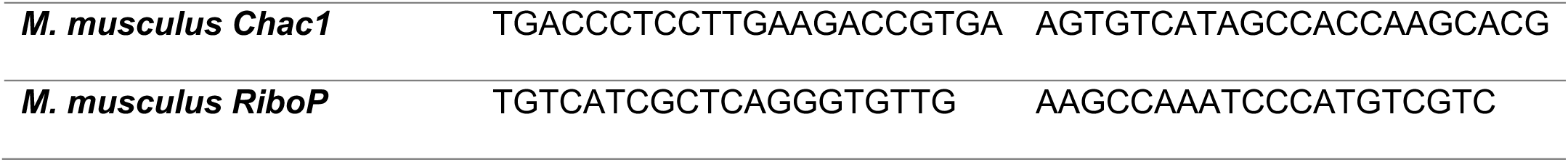

### Measurement of Mitochondrial Membrane Potential

Cells were seeded at a density of 200,000 cells per well in a 6-well plate and treated with 500 nM Tg for 3h prior to collection. CCCP (10 µM) was added 50 min before collection, followed by 200 nM TMRE (Thermofisher) 20 min before collection. Following treatments, cells were washed twice with phosphate-buffered saline (PBS) and dissociated using TrypLE Express (Thermo Fisher). The enzymatic reaction was neutralized through addition of flow buffer containing PBS and five percent fetal bovine serum (FBS). Fluorescence intensity of TMRE (552/574 nm) for 20,000 cells/condition was measured on a Bio-Rad ZE5 Cell Analyzer using the 561 nm green-yellow laser in combination with the 577/15 filter. Analysis was performed using FlowJo^TM^ Software (BD Biosciences). Data are presented as geometric mean of the fluorescence intensity from three experiments normalized to vehicle-treated cells.

### Cell Proliferation and Apoptosis Assays

*Perk^+/+^* and *Perk^-/-^* MEFs were treated for 6 h with either vehicle (DMSO) or the indicated drugs. Cells were washed with room temperature PBS and then trypsinized. Fresh media was added and cells were counted using a Countess Automated Cell Counter (ThermoFisher). Cells were seeded in a 6-well plate at three different dilutions: 30,000, 15,000 and 7,500 and then allowed to proliferate for five days while incubated at 37°C. At five days, media was aspirated off and cells were rinsed with PBS. Cells were then fixed with crystal violet staining solution (0.5% crystal violet (w/v), 20% methanol) for 10 minutes in room temperature. The crystal violet staining solution was then removed and cells were washed 3x with PBS and allowed to dry overnight. Images were then taken of the stained plates. Other parameters of ER Stress induced cell death were measured through immunoblotting or with the Caspase 3/7 Glo Assay Kit (Promega) according to the manufacturer’s protocol. In brief, cells were plated at 10k/well in a white 96-well plate. The next day, cells were treated with the indicated drugs for 24 h (n=5 for each condition). After treatment, an equal volume of Caspase 3/7 substrate was added to the wells and allowed to incubate at room temperature for 1 h at 37°C and 5% CO_2_. Luminescence was then measured in an Infinite F200 PRO plate reader (Tecan) with an integration time of 1000 ms.

### Mitochondrial Respiration

Mitochondrial respiration parameters were measured using a Mito Stress Test Kit and XF96 Extracellular Flux Analyzer (Seahorse Bioscience) according to the manufacturer’s protocol. In brief, 15,000 cells/well were plated on a 96-well plate on Cell-Tak (Corning) coated wells in their standard growing media and cultured overnight. The next day, cells were treated with drugs and incubated at 37°C, 5% CO_2_ for the indicated treatment time. Seahorse XF assay media (Agilent) was used to wash and remove the standard growing media from the cell plate and the sensor plate was prepped with standard Mito Stress Test drugs. After calibration of the sensor plate on the XF96 Extracellular Flux Analyzer, the cell plate was inserted into the machine and each parameter was calculated with 4 measurements separated by 5 min intervals following injections of drugs. Basal respiration measurement was followed by injections of oligomycin (2 µM), FCCP (0.5 µM) and Rotenone/Antimycin (1µM each), respectively. Individual respiratory parameters were then calculated according to the manufacturer’s protocol.

## ACKNOWLEDGEMENTS.

We thank Evan Powers for critical reading of this manuscript. This work was funded by the National Institutes of Health (NS095892, NS125674 to RLW), a National Science Foundation predoctoral fellowship (VP), and the George E. Hewitt Postdoctoral fellowship (VD).

## AUTHOR CONTRIBUTIONS

VP, KRB, and VD conceived, designed, performed, and interpreted the experiments. GA, JDR, XG performed experiments. JDR and RLW interpreted experiments. MK and RLW supervised the project. VP, KRB, and RLW wrote the manuscript. VP, KRB, VD, GA, JDR, XG, MK, and RLW provided critical revisions to the manuscript and approval for submission.

---

**Supplement to Fig. 1.**
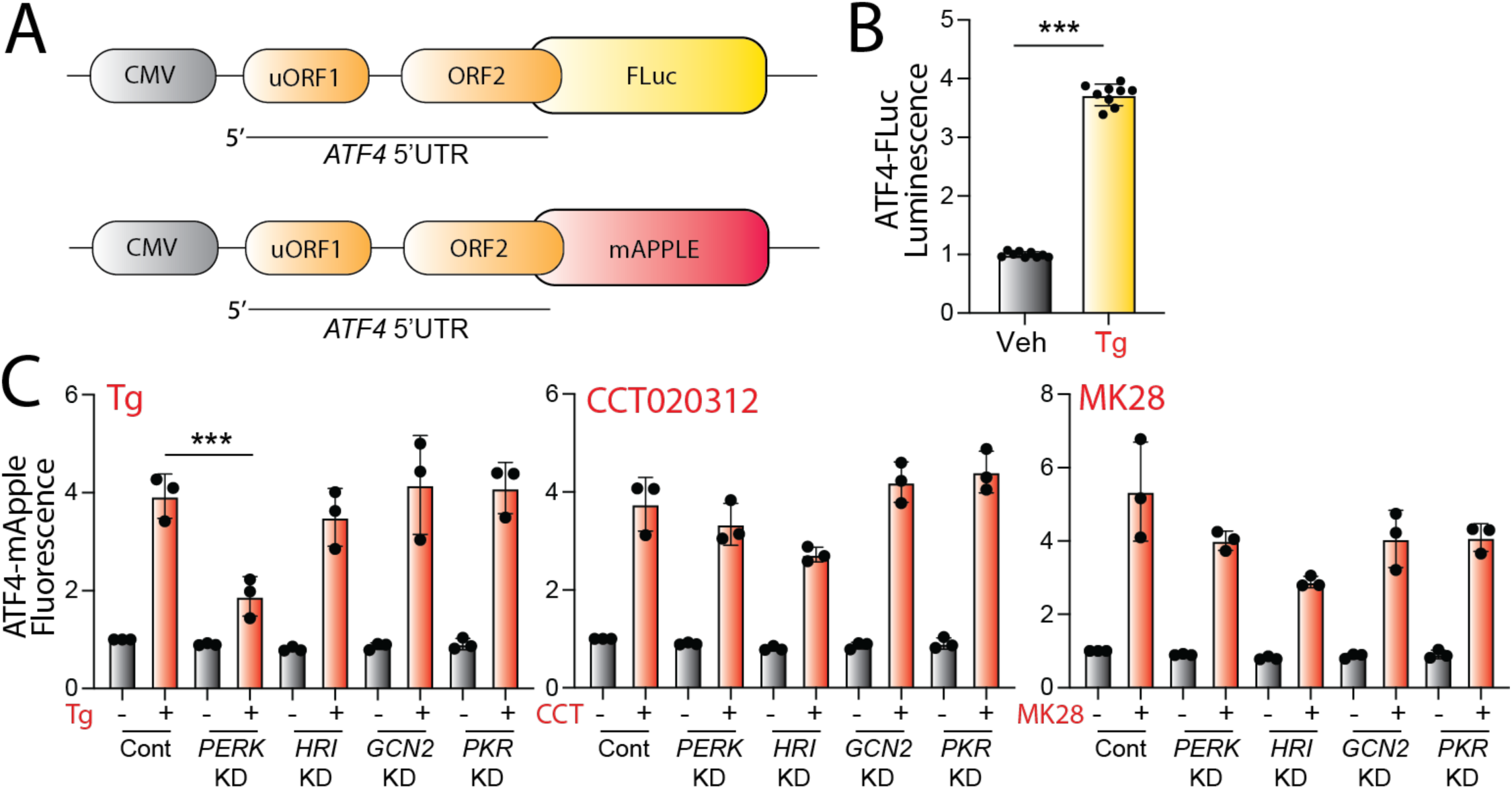
Pharmacologic activation of integrated stress response (ISR) kinases. **A.** Illustration showing the ATF4-FLuc and ATF4-mApple ISR reporters (Guo *et al*., 2020; Yang *et al*., 2022). **B.** Activation of ATF4-FLuc ISR reporter stably expressed in HEK293T cells treated for 3 h with thapsigargin (Tg; 500 nM). Error bars show SEM for n=9 replicates. **C**. Graphs showing activation of the ATF4-mApple ISR reporter stably expressed in HEK293T cells CRISPR-depleted of the indicated ISR kinase and treated for 8 h with BtdCPU (10 µM), halofuginone (100 nM), Erlotinib (25 µM), or Sunitinib (10 µM). Error bars show SEM for n=3 replicates. ***p<0.005 for one-way ANOVA.

**Supplement to Fig. 2.**
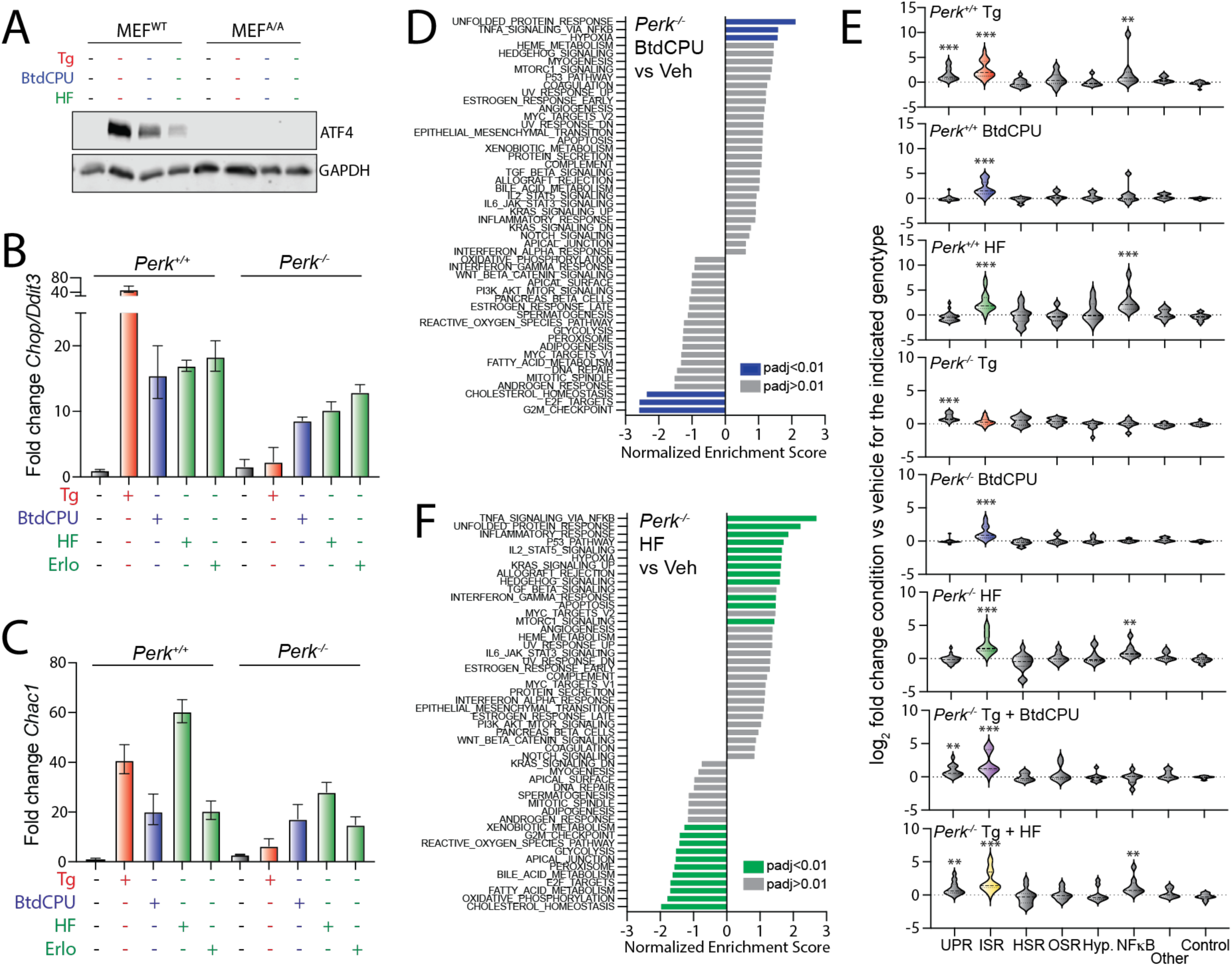
Pharmacologic ISR activators restore ISR signaling in *Perk*-deficient MEFs. **A.** Immunoblot of lysates prepared from MEF^WT^ and MEF^A/A^ cells treated for 3 h with thapsigargin (Tg; 500 nM), BtdCPU (10 µM), halofuginone (HF; 100 nM). **B,C.** Expression of the ISR targets *Chop/Ddit3* or *Chac1* in *Perk^+/+^* and *Perk^-/-^* MEFs treated for 6 h with thapsigargin (Tg, 500 nM), BtdCPU (10 µM), halofuginone (100 nM), or erlotinib (25 µM), as indicated. Error bars show 95% confidence interval. **D**. Gene set enrichment analysis (GSEA) for hallmark genesets of RNAseq data from *Perk^-/-^* MEFs treated for 6 h with BtdCPU (10 µM). Full GSEA is included in **Table S3**. **E**. Expression, measured by RNAseq, of genesets comprising target genes of the UPR (IRE1/ATF6), ISR, heat shock response (HSR), oxidative stress response (OSR), hypoxic stress response (Hyp.), NFκB inflammatory response, other stress-responsive genes, and control genes. Geneset are shown in **Table S4**. **F**. Gene set enrichment analysis (GSEA) for hallmark genesets of RNAseq data from *Perk^-/-^* MEFs treated for 6 h with halofuginone (HF, 100 nM). Full GSEA is included in **Table S3**.

**Supplement to Fig. 3.**
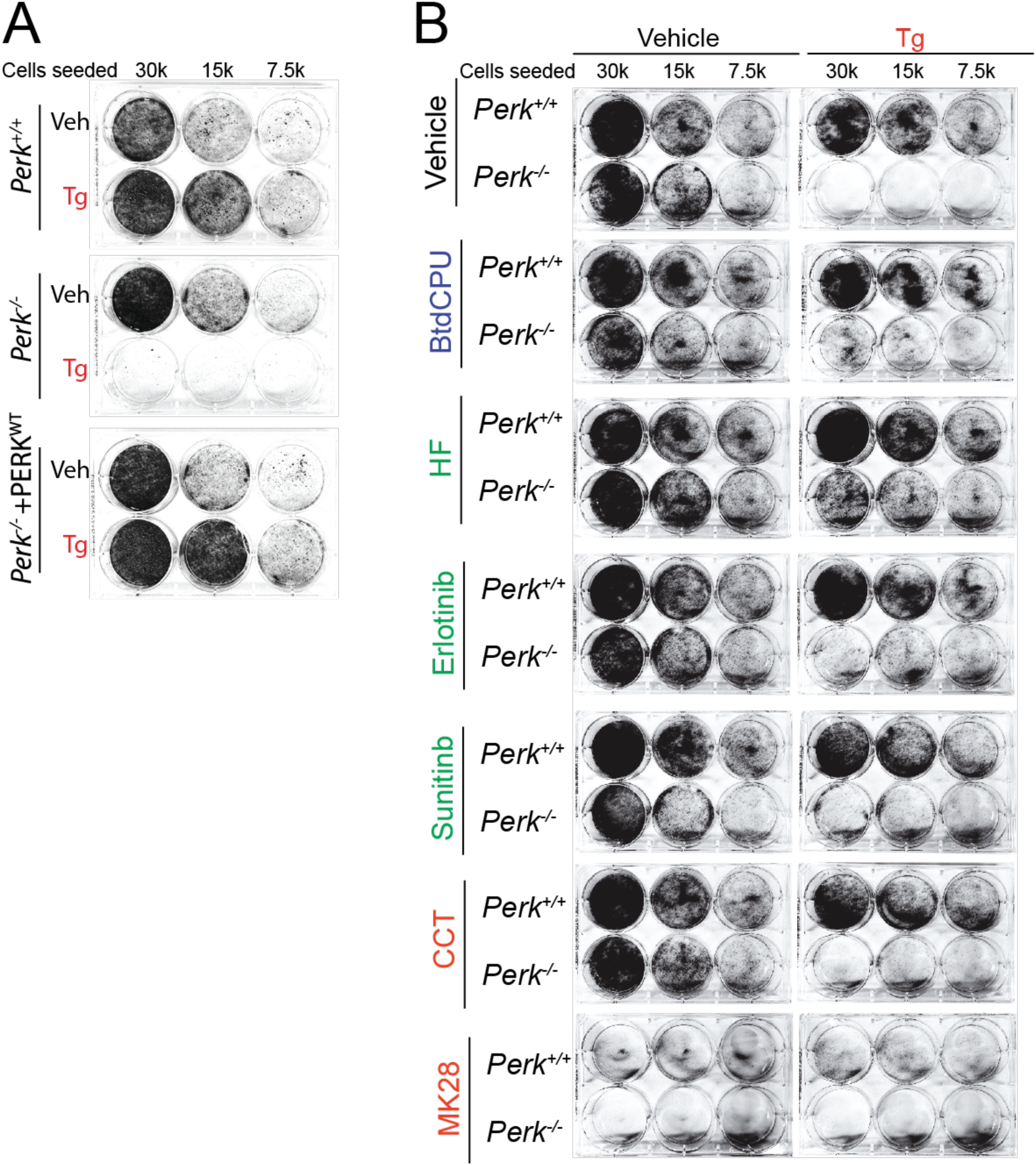
BtdCPU and halofuginone reestablish ER stress sensitivity in *Perk*-deficient MEFs. **A.** Representative crystal violet staining of *Perk^+/+^*, *Perk^-/-^* and *Perk^-/-^* MEFs with wild-type *Perk* overexpressed treated for 6 h with thapsigargin (Tg; 500 nM) and then replated and allowed to proliferate in 6 well plates. Crystal violet staining was performed 72 h after replating. **B**. Representative crystal violet staining of *Perk^+/+^* and *Perk^-/-^*MEFs treated for 6 h with thapsigargin (Tg; 500 nM) and or BtdCPU (10 µM), halofuginone (100 nM), Erlotinib (25 µM), Sunitinib (10 µM), CCT020312 (10 µM), and MK28 (20 µM). and then replated and allowed to proliferate in 6 well plates. Crystal violet staining was performed 72 h after replating.

**Supplement to Fig. 4.**
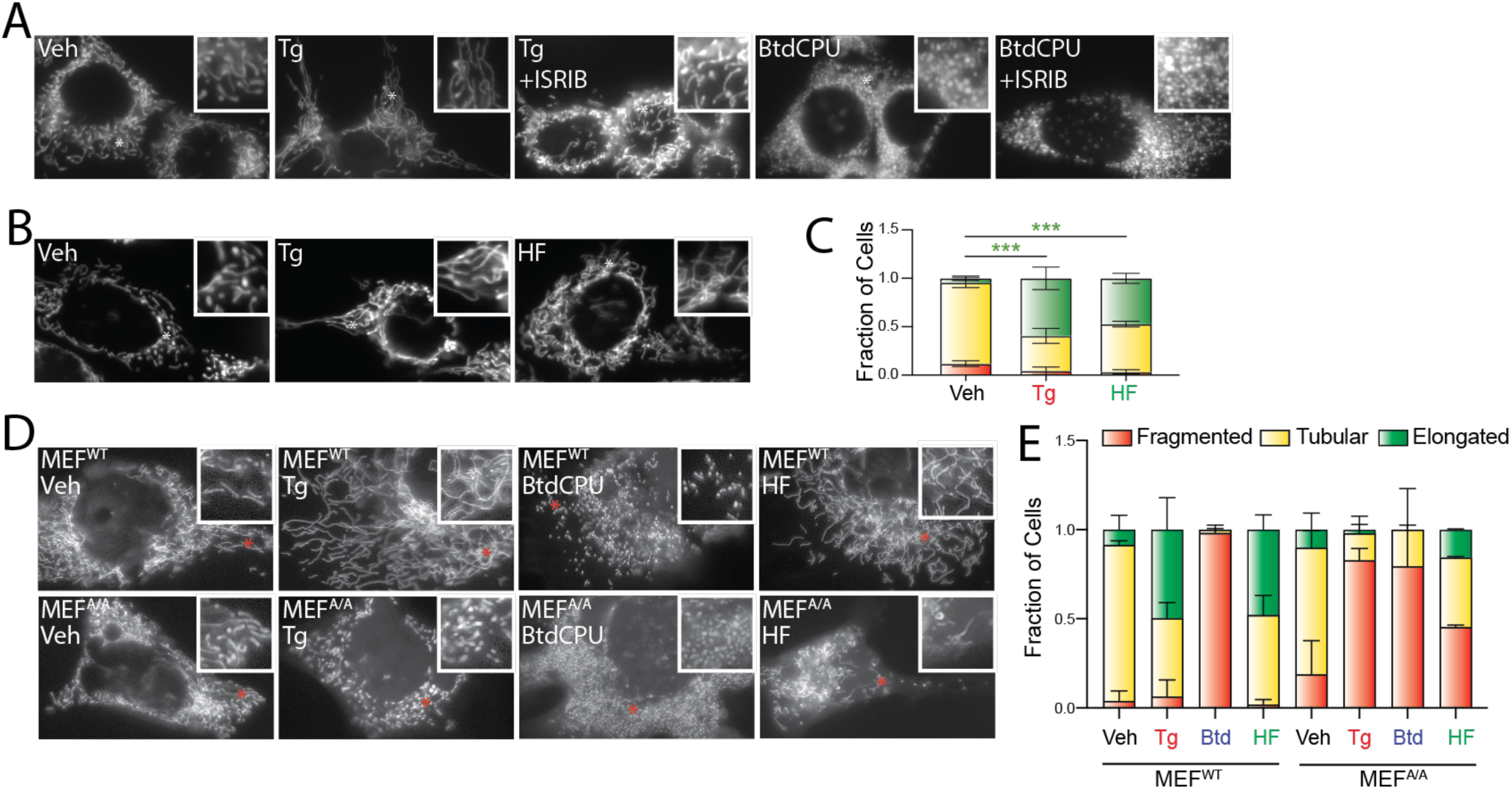
BtdCPU and halofuginone differentially impact mitochondrial morphology. **A.** Representative images of mitochondrial morphology in MEF cells stably expressing mitochondrial-targeted GFP treated for 3 h with thapsigargin (Tg; 500 nM), BtdCPU (10 µM), and/or ISRIB (200 nM). **B, C**. Representative images and quantification of mitochondrial morphology in MEF cells stably expressing mitochondrial-targeted GFP treated for 3 h with thapsigargin (Tg; 500 nM) or halofuginone (HF; 100 nM). Error bars show SEM for n=2 replicates. ***p<0.005 for two-way ANOVA. (green indicates comparisons between elongated mitochondria fractions). **D, E**. Representative images and quantification of mitochondrial morphology in MEF^WT^ and MEF^A/A^ cells expressing mitochondrial targeted GFP (^mt^GFP) and treated for 3 h with thapsigargin (Tg; 500 nM), BtdCPU (10 µM), halofuginone (HF; 100 nM). The inset shows 2-fold magnification of the image centered on the white asterisk. Error bars show SEM for n=2 replicates.

**Supplement to Fig. 5.**
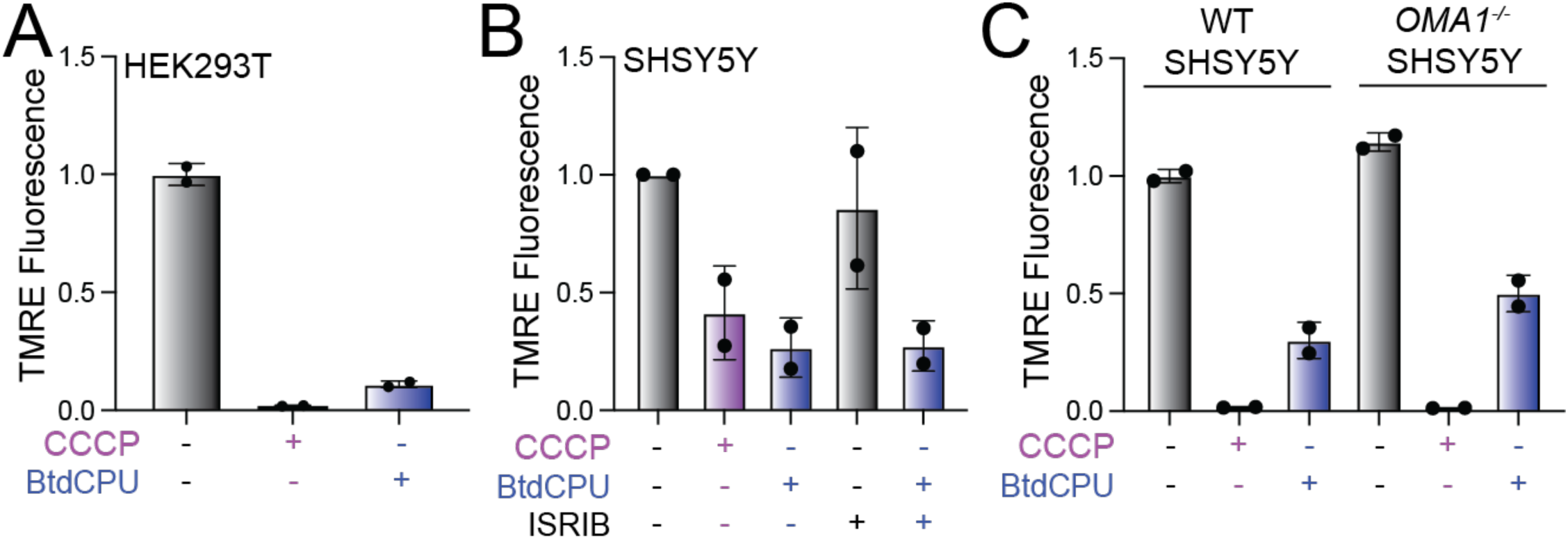
BtdCPU promotes mitochondrial uncoupling. **A**. Mitochondrial membrane potential, measured by TMRE fluorescence, in HEK293T cells pre-treated for 3 h with BtdCPU (10 µM) or for 30 min with CCCP (10 μM). **B**. Mitochondrial membrane potential, measured by TMRE fluorescence, in SHSY5Y cells pre-treated for 3 h with BtdCPU (10 µM) and/or ISRIB (200 nM) or for 30 min with CCCP (10 μM). **C**. Mitochondrial membrane potential, measured by TMRE fluorescence, in SHSY5Y cells or *OMA1* knockout SHSY5Y cells pre-treated for 3 h with BtdCPU (10 µM) or for 30 min with CCCP (10 μM).

**Supplement to Fig. 6.**
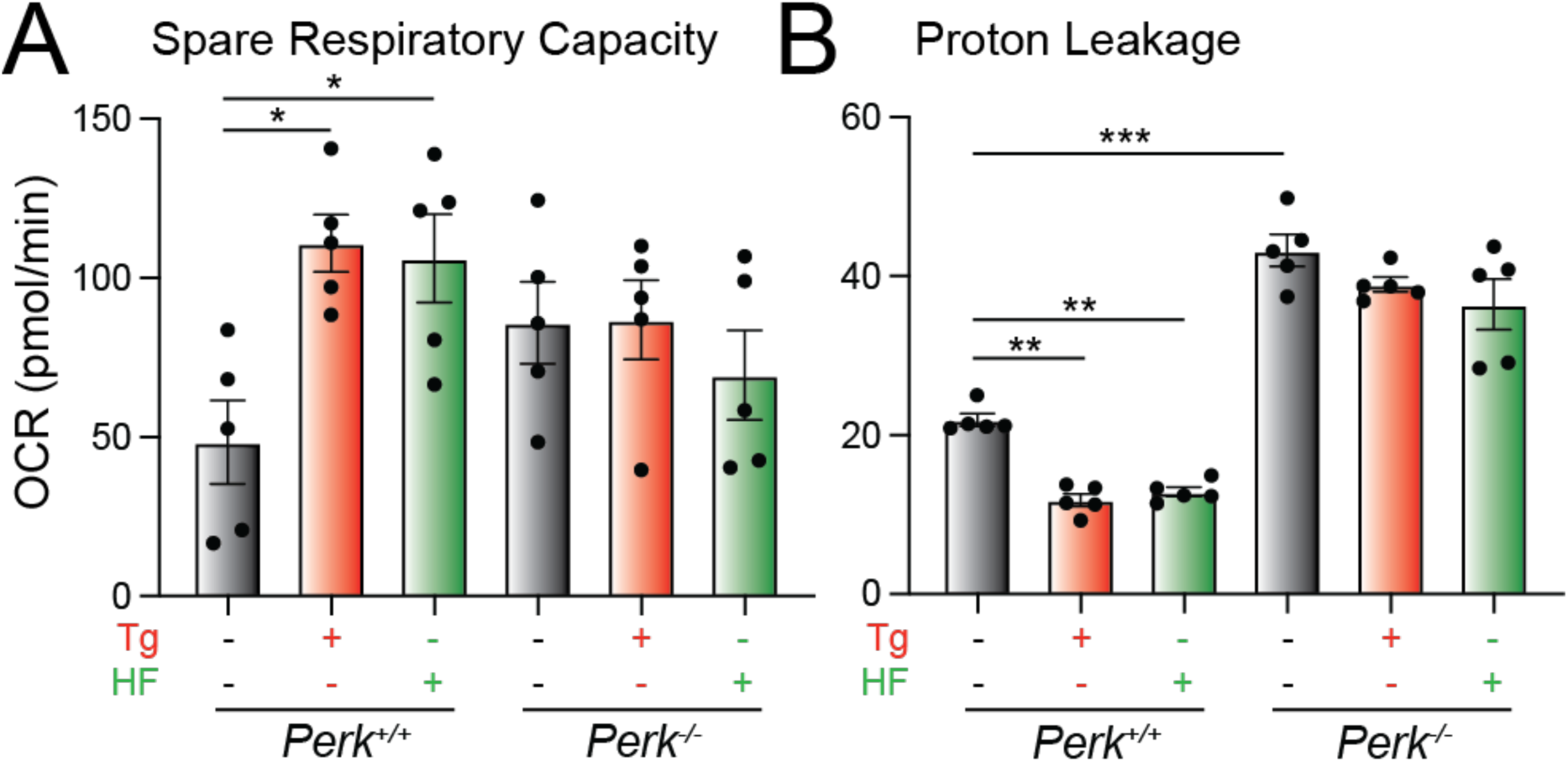
Halofuginone promotes adaptive remodeling of mitochondria respiration in PERK-deficient cells. **A,B.** Spare respiratory capacity (**A**) and proton leak (**B**) in *Perk^+/+^* and *Perk^-/-^* MEFs treated for 3 h with thapsigargin (Tg; 500 nM) or halofuginone (HF, 100 nM) measured from the mitochondrial stress test shown in Fig. 6C. Error bars show SEM for n=5 replicates. *p<0.05, **p<0.01, ***p<0.005 from one-way ANOVA.

## SUPPLEMENTAL TABLE LEGENDS

**Table S1. (Supplement to Figure 2 and Figure S2. Pharmacologic ISR activators restore ISR signaling in *Perk*-deficient MEFs).** DESEQ2 from RNAseq of *Perk^+/+^* and *Perk^-/-^* MEFs treated for 6 h with thapsigargin (Tg; 500 nM), BtdCPU (10 µM), and/or halofuginone (HF; 100 nM). The complete RNA-seq data is deposited in gene expression omnibus (GEO) as GSE227134.

**Table S2. (Supplement to Figure 2 and Figure S2. Pharmacologic ISR activators restore ISR signaling in *Perk*-deficient MEFs).** Expression, measured by RNAseq, of unfolded protein response (UPR; IRE1/XBP1s and ATF6) target genes and integrated stress response (ISR) target genes, as defined in Grandjean et al (2019) *ACS Chem Biol*, in *Perk^+/+^*and *Perk^-/-^* MEFs treated for 6 h with thapsigargin (Tg; 500 nM), BtdCPU (10 µM), and/or halofuginone (HF; 100 nM). The expression of individual genes is shown normalized to that observed in Tg-treated *Perk^+/+^* MEFs, as described in Grandjean et al (2019) *ACS Chem Biol*.

**Table S3. (Supplement to Figure 2 and Figure S2. Pharmacologic ISR activators restore ISR signaling in *Perk*-deficient MEFs).** Geneset Enrichment analysis (GSEA) of RNAseq data from *Perk^+/+^* and *Perk^-/-^*MEFs treated for 6 h with thapsigargin (Tg; 500 nM), BtdCPU (10 µM), and/or halofuginone (HF; 100 nM). There are 6 individual tabs in this workbook describing *h^+/+^* MEF treated with Tg, *Perk^+/+^* MEFs treated with BtdCPU, *Perk^+/+^* MEFs treated with HF, *Perk^-/-^* MEFs treated with Tg, *Perk^-/^*MEFs treated with BtdCPU, and *Perk^-/^* MEFs treated with HF.

**Table S4. (Supplement to Figure 2 and Figure S2. Pharmacologic ISR activators restore ISR signaling in *Perk*-deficient MEFs).** Expression, measured by RNAseq, of genes regulated by stress-responsive signaling pathways including the unfolded protein response (UPR), integrated stress response (ISR), heat shock response (HSR), oxidative stress response (OSR), and NFκB inflammatory response in *Perk^+/+^* and *Perk^-/-^* MEFs treated for 6 h with thapsigargin (Tg; 500 nM), BtdCPU (10 µM), and/or halofuginone (HF; 100 nM).

